# A miR-124-mediated post-transcriptional mechanism controlling the cell fate switch of astrocytes to induced-neurons

**DOI:** 10.1101/2020.06.01.127126

**Authors:** Elsa Papadimitriou, Paraskevi N. Koutsoudaki, Irini Thanou, Timokratis Karamitros, Dimitra Karagkouni, Dafni Chroni-Tzartou, Maria Gaitanou, Christos Gkemisis, Maria Margariti, Evangelia Xingi, Socrates J. Tzartos, Artemis G. Hatzigeorgiou, Dimitra Thomaidou

**Affiliations:** Neural Stem Cells and Neuroimaging Group, Department of Neurobiology, Hellenic Pasteur Institute; Bioinformatics and Applied Genomics Unit, Department of Microbiology, Hellenic Pasteur Institute; DIANA-Lab, Hellenic Pasteur Institute & Dept. of Computer Science and Biomedical Informatics, Univ. of Thessaly; Laboratory of Molecular Neurobiology and Immunology, Department of Neurobiology, Hellenic Pasteur Institute; Laboratory of Cellular and Molecular Neurobiology – Stem Cells, Department of Neurobiology, Hellenic Pasteur Institute; Light Microscopy Unit, Hellenic Pasteur Institute

**Keywords:** astrocytes, direct reprogramming, ISX9, miR-124, Zfp36l1

## Abstract

The miRNA miR-124 has been employed supplementary to neurogenic TFs and other miRNAs to enhance direct neurogenic conversion by suppressing multiple non-neuronal targets. Aim of the study was to investigate whether miR-124 is sufficient to drive direct reprogramming of astrocytes to induced-neurons (iNs) on its own and elucidate its independent mechanism of reprogramming action. Our data show that miR-124 is a potent driver of the reprogramming switch of astrocytes towards an immature neuronal fate, by directly targeting the RNA-binding protein Zfp36l1 implicated in ARE-mediated mRNA decay and subsequently de-repressing Zfp36l1 neurogenic interactome. To this end miR-124 contribution in iNs’ production largely recapitulates endogenous neurogenesis pathways, being further enhanced upon addition of the neurogenic compound ISX9, which greatly improves both miR-124-induced reprogramming efficiency and iNs’ functional maturation. Importantly, miR-124 is potent to guide direct conversion of reactive astrocytes to immature iNs of cortical identity *in vivo* following cortical trauma, confirming its ‘master’ reprogramming capacity within the injured cortical microenvironment, while ISX9 supplementation confers a survival advantage to newly produced iNs.

## Introduction

Direct astrocytic reprogramming to induced-neurons (iNs) is a powerful approach for manipulating cell fate, as it takes advantage of the intrinsic neural stem cell (NSC) potential of reactive astrocytes (Magnusson et al. 2014), while it offers the possibility of reprogramming resident brain cells. To this end astrocytic cell fate conversion to iNs has been well-established *in vitro* (Berninger et al. 2007; Heinrich et al. 2010; Aravantinou-Fatorou et al. 2015)and *in vivo* (Guo et al. 2014; Torper et al. 2013; Mattugini et al. 2019) using combinations of transcription factors (TFs) or chemical cocktails (L. Zhang et al. 2015; Li et al. 2015; L. Gao et al. 2017). Challenging the expression of lineage-specific TFs during reprogramming is accompanied by changes in the expression of regulatory RNAs, mostly miRNAs, that post-transcriptionally modulate high numbers of neurogenesis-promoting factors and to this end miRNAs have been introduced, supplementary or alternatively to TFs, to instruct direct neuronal reprogramming (Yoo et al. 2011).

Among neurogenic miRNAs, miR-124 has been shown to contribute to efficient neurogenic conversion of fibroblasts when coupled with certain TFs or other miRNAs, in particular miR-9/9* and the strong neuronal reprogramming potential of this cocktail has been elaborately studied at the transcriptomic and epigenetic level (Ambasudhan et al. 2011; Abernathy et al. 2017; Wohl and Reh 2016; Victor et al. 2014). miR-124 acts globally to increase the expression levels of neuronal genes by repressing components of major neuronal gene repressor complexes, such as the anti-neural transcriptional repressor REST complex (Visvanathan et al. 2007; Baudet et al. 2012; Volvert et al. 2014) and the Polycomb Repressive Complex 2 (PRC2) (Neo et al. 2014; Lee et al. 2018), while it also participates at the post-transcriptional regulation of neuronal transcripts by targeting the neuron-specific splicing global repressor Ptbp1 (Makeyev et al. 2007). Besides its roles in transcriptional and post-transcriptional regulation, miR-124 is a key mediator of a chromatin permissive environment for neuronal reprogramming through its involvement in the formation of the neuron specific chromatin remodeling complex nBAF (Yoo et al. 2009).

However, although miR-124 has been lately utilized in many reprogramming cocktails for the neurogenic conversion of fibroblasts (Birtele et al. 2019; Jiang et al. 2015; Victor et al. 2018; Ambasudhan et al. 2011; Yoo et al. 2011), neither its potential to induce fate conversion of astrocytes to induced-neurons (iNs) *in vitro* or *in vivo*, nor its mechanism of action in instructing direct reprogramming on its own have been investigated. In this study we show that miR-124 is sufficient to instruct reprogramming of cortical astrocytes to immature iNs *in vitro* controlling the reprogramming “switch” of astrocytes towards the neuronal fate by down-regulating genes with important regulatory roles in astrocytic function. Among these we identified for the first time the RNA binding protein Zfp36l1, implicated in ARE-mediated mRNA decay (Lai et al. 2000), as a direct target of miR-124 and further found certain neuronal-specific Zfp36l1 targets that participate in cortical development being de-repressed in miR-124-iNs. Importantly, by blocking miR-124 specific binding in Zfp36l1 3’UTR, we revealed that miR-124/Zfp36l1 interaction is one of the drivers of miR-124-induced astrocytic fate switch and induction of neuronal identity. To enhance the neuronal differentiation of reprogrammed immature iNs, we combined miR-124 with isoexasole-9 (ISX9) chemical compound known to possess neurogenesis-promoting properties (Schneider et al. 2008; Li et al. 2015). Functional analysis of the two molecules’ combination revealed that *in vitro* addition of ISX9 promoted both the neurogenic conversion and greatly enhanced the functional maturation of miR-124+ISX9-iNs. Importantly, *in vivo* miR-124 was also potent either alone or along with ISX9, to guide neuronal reprogramming of reactive astrocytes following cortical trauma to immature iNs, present after 8 weeks, with ISX9 contributing to iNs’ enhanced survival, revealing the *in vivo* ‘master’ reprogramming capacity of miR-124 within the injured cortical micro-environment.

## Results

### miR-124 is sufficient to instruct reprogramming of postnatal cortical astrocytes to immature induced-neurons

To study the potential of miR-124 to instruct neuronal reprogramming of astrocytes on its own, cultured postnatal day3-5 (P3-5) mouse cortical astrocytes were transfected with miR-124-3p mimics (**Fig 1A**). Firstly, we verified that the initial primary astrocytic culture comprised majorly of GFAP+ astrocytes (72.2% ± 8.6%), while only 2.3%± 2.1%were Tuj1+ neurons (**Suppl Fig 1A and B**). BrdU administration for the first 4 days of reprogramming revealed that about 70% of cells at day7 (d7) (astrocytes treated or not with scrambled miRNA, sc-miRNA) had undergone cell division, while overexpression of miR-124 significantly reduced this percentage to 50% (**Suppl Fig 1D and E**). After 1 week (d7) nearly 35% of miR-124-treated cells were Tuj1+, exhibiting multipolar morphology (**Fig 1B and C**), as compared to control astrocytes that received sc-miRNA, where no Tuj1positivity was detected (**Suppl Fig 1C**). Still, miR-124-iNs exhibited low differentiation potential and only 19% of the cells remained Tuj1+ at d14 of the reprogramming process (**Fig 1B and C**). Notably almost 60% of Tuj1+ miR-124-iNs at d7 had incorporated BrdU (**Supl Fig 1F**), accounting for nearly 20% of the total cells in culture (**Suppl Fig 1G**), indicating their non-neuronal origin. The ability of miR-124 to instruct neurogenic reprogramming was further supported by RT-qPCR expression analysis of several neurogenic transcription factors (TFs) at d7, where miR-124 overexpression induced the up-regulation of the mRNA levels of the proneural TFs *Mash1* and to a lesser extend *Neurog2* (**Fig 1D**), while it additionally up-regulated TFs related to both dorsal (*Tbr2, Tbr1, Fezf2* and *Cux1*) (**Fig 1E**) and ventral telencephalon development (*Gsx2, Dlx1*) (**Fig 1F**). We also observed up-regulation of TFs related to neuronal differentiation (*Sox4, Sox11, Hes6*) (**Fig 1G**), however we failed to detect an up-regulation of *NeuroD1* (**Fig 1G**), which is known to play a crucial role in neuronal reprogramming (Pataskar et al. 2016; Matsuda et al. 2019; Guo et al. 2014). Instead, miR-124 significantly reduced *NeuroD1* mRNA levels, implying that this reduction may contribute to the low differentiation capacity of miR-124-iNs. Further, immunofluorescence analysis indicated that the majority of miR-124-iNs (nearly 80%) were Mash1+ and also exhibited low Tbr2 expression (nearly 70%), while only a small percentage of reprogrammed cells (15%) were positive for the ventral TF Gsx2 (**Fig 1H and I**), indicating that the Mash1-Tbr2 trajectory is most prominently activated in the majority of miR-124-iNs.

**Figure 1:**
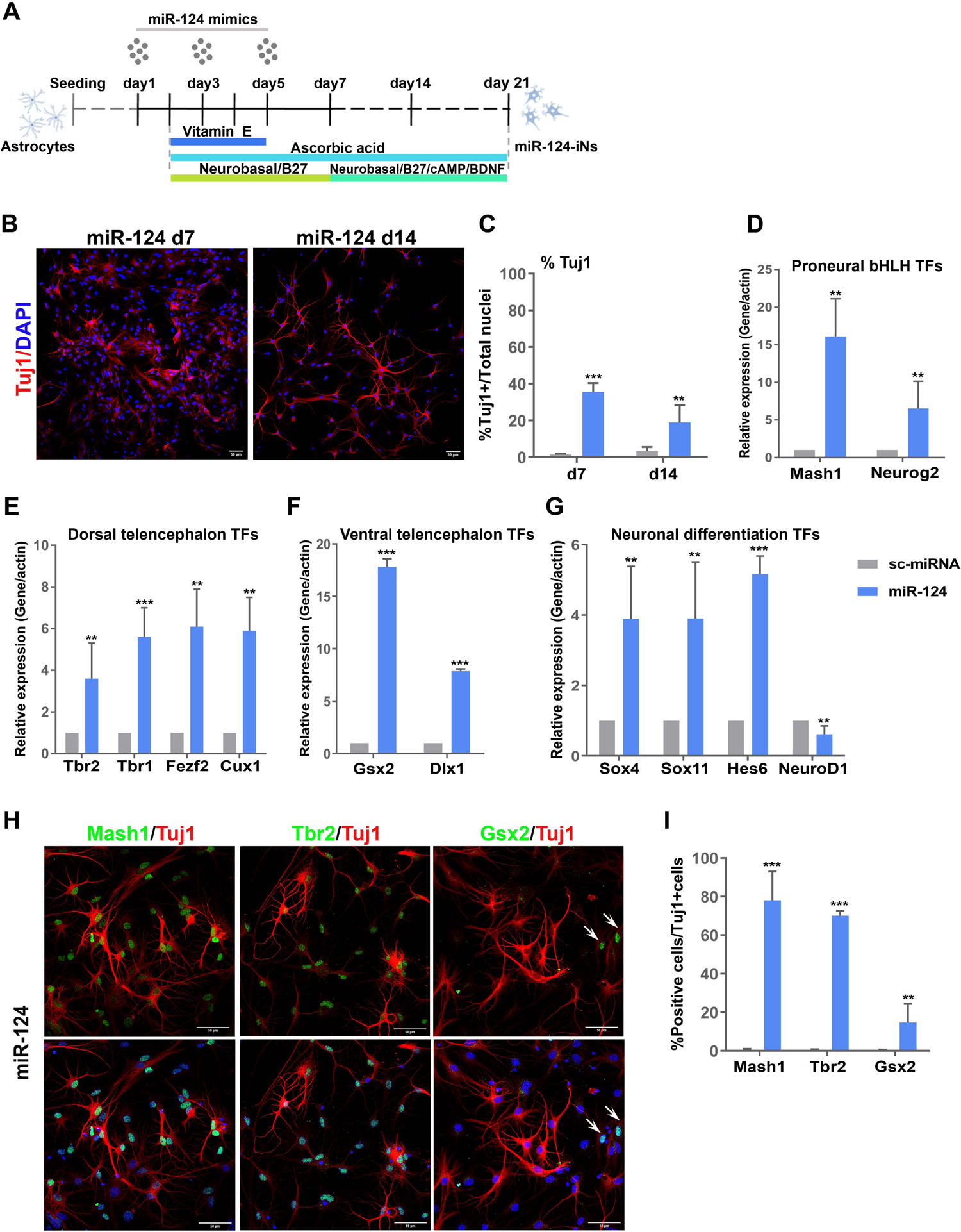
miR-124 is sufficient to instruct reprogramming of postnatal cortical astrocytes to iNs. **(A)** Overview of the miR-124-mediated reprogramming protocol. **(B)** Immunostaining of astrocytes reprogrammed with miR-124 at d7 and d14 of the reprogramming protocol with anti-Tuj1 antibody. **(C)** Quantification of the percentage of Tuj1+ reprogrammed cells (average ±SD, n=4 independent experiments). RT-qPCR analysis of the mRNA levels of the proneural TFs, *Mash1* and *Neurog2* **(D)**, the dorsal telencephalon TFs, *Tbr2, Tbr1, Fezf2, Cux1* **(E)**, the ventral telencephalon *TFs, Gsx2* and *Dlx1* **(F),** and the neuronal differentiation TFs, *Sox4, Sox11, Hes6* and *NeuroD1* **(G)**. Data are presented as fold change *vs* sc-miRNA (average ± SD, n=3 independent experiments). **(H)** Co-immunostaining of astrocytes reprogrammed with miR-124 at d7 of the reprogramming protocol with anti-Mash1/Tuj1, anti-Tbr2/Tuj1 and anti-Gsx2/Tuj1 antibodies. **(I)** Quantification of the percentage of Mash1+, Tbr2+ and Gsx2+ in Tuj1+ reprogrammed cells (average ± SD, n=3 independent experiments). For all presented data **p<0.01 and ***p<0.001 vs sc-miRNA.

### The neurogenic compound ISX9 greatly enhances the miR-124-induced reprogramming efficiency and differentiation state of iNs

The observed down-regulation of NeuroD1 by miR-124 prompted us to supplement the reprogramming medium from d2 to d10 with the chemical compound ISX9, known to up-regulate NeuroD1 levels and enhance neuronal differentiation (Schneider et al. 2008). Indeed, ISX9 addition led to the acquisition of a more differentiated neuronal phenotype with a smaller soma and longer projections (**Fig 2A**) and significantly increased the percentage of Tuj1+ iNs from 35% to 62% at d7 and from 19% to 38% at d14 (**Fig 2B**). Importantly, ISX9 was potent to reverse the miR-124-induced reduction of *NeuroD1* mRNA levels (**Fig 2C**) and to greatly elevate the transcriptional levels of *Neurog2* and *Tbr2* peaking at d7 (**Fig 2D**), while it also induced a moderate reduction in *Mash1* mRNA levels (**Fig 2E**). Importantly, ISX9 was not able to induce reprogramming of astrocytes on its own (sc-miRNA+ISX9) (**Suppl Fig 2A**), despite evoking robust up-regulation of the mRNA levels of *NeuroD1* (**Fig 2D**) and other neurogenic TFs (**Suppl Fig 2B**) and to a small, but significant, extend the protein levels of Mash1 and Tbr2 (**Suppl Fig 2C and D**). Additionally, supplementation of ISX9 along with miR-124 significantly increased the percentage of Tbr2+/Tuj1+ iNs, without affecting the percentages of Mash1+/Tuj1+ iNs or Gsx2+/Tuj1+ iNs relatively to miR-124 alone (**Fig 2F and G**). Still, measurement of the mean nuclear fluorescence intensity per cell in miR-124-iNs and miR-124+ISX9-iNs at d7, revealed a significant enhancement of Tbr2 protein levels (**Fig 2H**) and a down-regulation of Mash1 levels (**Fig 2I**) following ISX9 addition.

**Figure 2:**
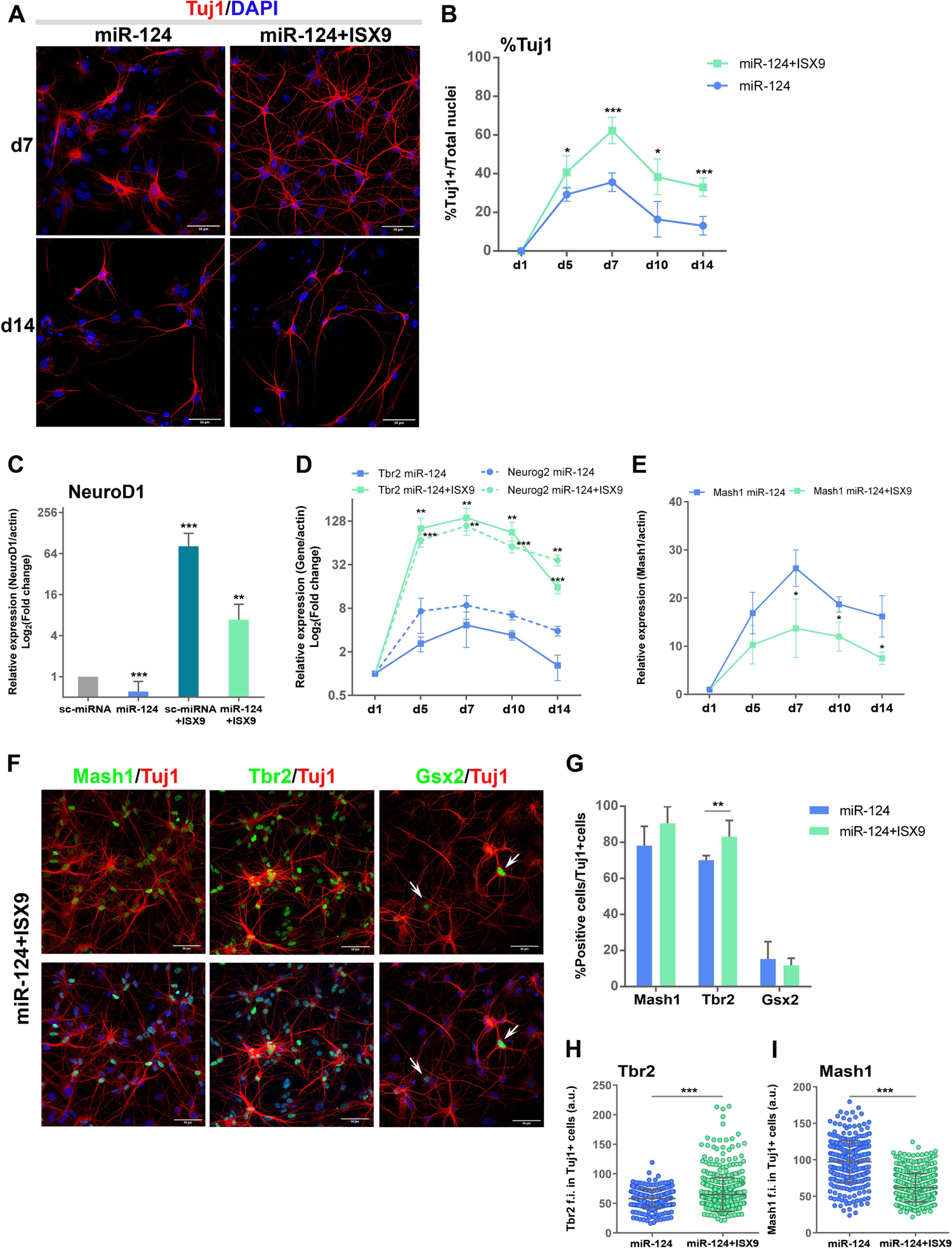
The neurogenic compound ISX9 greatly enhances the miR-124-induced reprogramming efficiency and differentiation state of iNs. **(A)** Immunostaining of astrocytes reprogrammed with miR-124 or miR-124+ISX9 at d7 and d14 of the reprogramming protocol with anti-Tuj1 antibody. **(B)** Quantification of the percentage of Tuj1+ reprogrammed cells at the time points d1, d5, d7, d10 and d14 of the reprogramming protocol (average ± SD, n=3 independent experiments for d5 and d10, n=8 for d7 and n=4 for d14, *p<0.05 and ***p<0.001 *vs* miR-124). **(C)** RT-qPCR analysis of the mRNA levels of NeuroD1 at d7 of the reprogramming protocol. Data are presented as log_2_ (fold change) *vs* sc-miRNA (average ± SD, n=3 independent experiments, **p<0.01 and ***p<0.001 *vs* sc-miRNA). RT-qPCR analysis of the mRNA levels of the TFs, Neurog2 and Tbr2 **(D)** and Mash1 **(E)** at the time points d1, d5, d7, d10 and d14 of the reprogramming protocol. Data are presented as log_2_(fold change) (**D**) and fold change (**E**) *vs* astrocytes (d1) (average ± SD, n=3 independent experiments, *p<0.05, **p<0.01 and ***p<0.001 *vs* miR-124). **(F)** Co-immunostaining of astrocytes reprogrammed with miR-124+ISX9 at d7 of the reprogramming protocol with anti-Mash1/Tuj1, anti-Tbr2/Tuj1 and anti-Gsx2/Tuj1 antibodies. **(G)** Quantification of the percentage of Mash1+, Tbr2+ and Gsx2+ in Tuj1+ reprogrammed cells either with miR-124 or miR-124+ISX9 at d7 (average ± SD, n=3 independent experiments, **p<0.01 *vs* miR-124). Measurement of the mean nuclear fluorescence intensity of Tbr2 **(H)** and Mash1 **(I)** in Tuj1+ reprogrammed cells either with miR-124 or miR-124+ISX9 at d7 (co-immunostaining with anti-Tbr2/Mash1/Tuj1 antibodies). A representative experiment is shown of n=3 independent experiments (mean ± SD, n=326 cells for miR-124 and n=540 cells for miR-124+ISX9, ***p<0.001 *vs* miR-124).

These results led us to the inference that addition of ISX9 in the reprogramming medium reinforces the passage of miR-124+ISX9-iNs through a Tbr2+ intermediate stage, which seems to be the main reprogramming route these iNs follow. Interestingly, the addition of ISX9 also significantly enhanced the expression of *Insm1* (**Suppl Fig 2E**), a key transcriptional regulator of intermediate progenitors (IPs) (Elsen et al. 2018), further supporting the notion that iNs pass through an intermediate stage bearing molecular characteristics of endogenous IPs.

### miR-124+ISX9-iNs exhibit characteristics of mature, electrophysiologically active neurons

The majority of miR-124-iNs and miR-124+ISX9-iNs were positive for the cortical TF Tbr1 at d14 (**Suppl Fig 3A and B**). We mainly observed a moderate nuclear Tbr1 expression in miR-124-iNs, whereas miR-124+ISX9-iNs exhibited strong cytoplasmic Tbr1 expression, besides a moderate nuclear one (**Suppl Fig 3A**), a finding that has been previously reported in cortical neurons (Hong and Hsueh 2007). After 21d in culture, nearly 80% of Tuj1+ miR-124+ISX9-iNs were also positive for the mature neuronal markers MAP2 and Synapsin1, exhibiting a differentiated neuronal morphology (**Fig 3A and B**), while miR-124-iNs at d21 did not exhibit signs of further maturation and only a small percentage of them were MAP2+ and Synapsin1+ (**Fig 3B**). Additionally, at d28 the majority of miR-124+ISX9-iNs (nearly 90%) were positive for the glutamatergic marker vGlut1 (**Fig 3C and D**), while only 12% of them were positive for GABA (**Fig 3D and Suppl Fig 3C**).

**Figure 3:**
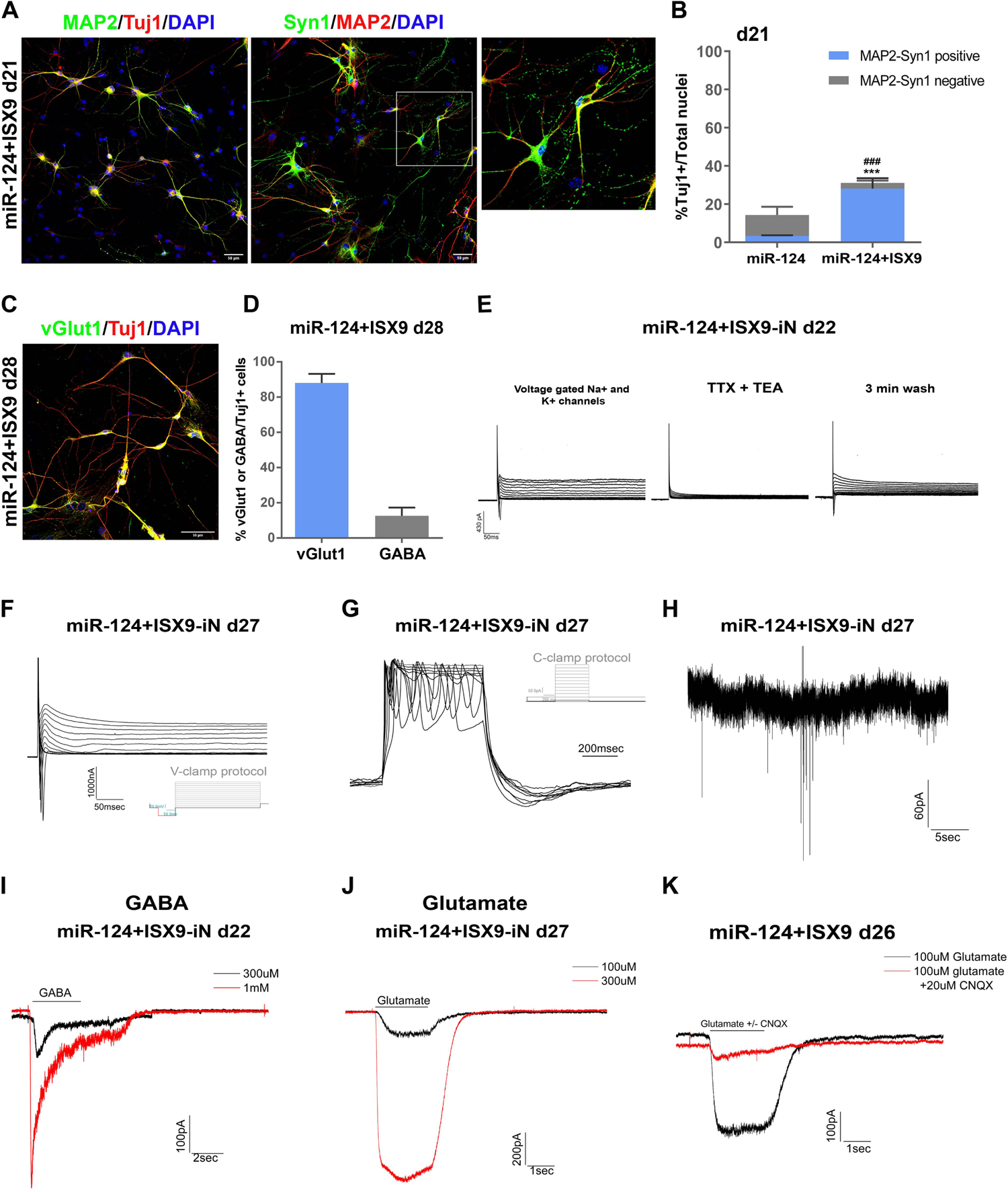
miR-124+ISX9-iNs exhibit characteristics of mature, electrophysiologically active neurons. **(A)** Co-immunostaining of miR-124+ISX9-iNs at d21 of the reprogramming protocol with anti-MAP2/Tuj1 and anti-MAP2/Synapsin1 antibodies. Inset area indicated in white frame. **(B)** Quantification of the percentage of Tuj1+ miR-124-iNs and miR-124+ISX9-iNs at d21 of the reprogramming protocol. The percentage of MAP2/Syn1 double positive (DP) Tuj1+ iNs is shown in blue (average ± SD, n=3 independent experiments, ***p<0.001 refers to %MAP2/Syn1 DP in Tuj1+ miR-124+ISX9-iNs *vs* miR-124-iNs and ^###^p<0.001 refers to %Tuj1 miR-124+ISX9-iNs *vs* miR-124-iNs). **(C)** Co-immunostaining of miR-124+ISX9-iNs at d28 of the reprogramming protocol with anti-vGlut1/Tuj1 antibodies. **(D)** Quantification of the percentage of vGlut1+/Tuj1+ and GABA+/Tuj1+ miR-124+ISX9-iNs at d28 (average ± SD, n=3 independent experiments). **(E)** Superimposed traces of inward Na^+^ currents and outward K^+^ currents evoked by depolarizing voltage steps obtained from a miR-124+ISX9-iN (d22) (left panel). Superimposed traces of inward and outward currents evoked by the same protocol after 1 min pre-application of 1μΜ TTX + 10mM TEA, showing the inhibition of both inward and outward currents (middle panel), followed by full recovery of the current traces after 3 min wash of the cell (right panel). **(F)** Superimposed traces of inward Na^+^ currents and outward K^+^ currents evoked by depolarizing voltage steps obtained from a miR-124+ISX9-iN (d27). **(G)** Example of a repetitive action potential induced from a mature miR-124+ISX9-iN (d27) by different current steps (injections) at the current clamp mode (the protocol of current clamp is shown upper right). (**H**) Example of a mature miR-124+ISX9-iN (d27) that exhibits spontaneous post-synaptic activity at −70 mV holding potential. Representative traces of ionic currents induced by application of the neurotransmitter GABA in two different concentrations (300μΜ and 1mM) obtained from a miR-124+ISX9-iN in the early stage of neuronal maturation (d22) **(I)** and the neurotransmitter glutamate in two different concentrations (100μΜ and 300μΜ) obtained from a miR-124+ISX9-iN in the late stage of maturation (d27) **(J)**.The cell membrane potential was held at −70 mV and the time of agonist application is indicated in bars above the traces. **(K)** Superimposed traces obtained from a mature miR-124+ISX9-iN (d26) with application of 100 μM glutamate (Glut) or co-application of 100μM Glut+CNQX indicated that the antagonist CNQX inhibits the AMPA/ kainite glutamate receptor.

To further establish the functional maturation of miR-124-iNs and miR-124+ISX9-iNs we performed electrophysiological analysis with whole-cell voltage-clamp and current-clamp recordings at different time points from d15 to d27. Rapidly inactivating inward Na^+^ currents and persistent outward K^+^ currents were recorded in miR-124+ISX9-iNs after d22 (n=47 out of 80 analyzed cells) in response to depolarizing voltage steps (**Fig 3E, left panel, Fig 3F**), while further application of TTX and TEA – selective Na^+^ channels’ (Na_V_) and K^+^ channels’ (K_V_) blockers respectively – confirmed that the Na_V_ were responsible for the inward currents and K_V_ for the outward currents (**Fig 3E, middle and right panels**). A small amount of K_V_ channels appeared in miR-124+ISX9-iNs before d21, while they became more evident in d23-27 (**Fig 3F**), in accordance with the ability of almost all recorded miR-124+ISX9-iNs (n=21 out of 30 recorded cells) to generate repetitive action potentials (APs) upon membrane depolarization (**Fig 3G**). Finally, rare spontaneous post-synaptic current activity was detected in few mature miR-124+ISX9-iNs (d27) (**Fig 3H**).On the other hand, miR-124-iNs exhibited lower amounts of Na_v_ and K_v_ channels and were thus not capable of firing action potentials (APs) (n=15cells).

The majority (80%) of miR-124+ISX9-iNs (**Fig 3I**) were capable of responding to different concentrations of GABA early in the course of their maturation (d22), even before the appearance of APs, which is in compliance with the expression of GABA receptors in early stages of neuronal development (Luján, Shigemoto, and López-Bendito 2005). Additionally, miR-124+ISX9-iNs were capable of responding to L-glutamate in a concentration dependent manner (**Fig 3J**), while L-glutamate-sensitive inward current was completely blocked after co-application of 100μΜ L-glutamate and 20μΜ CNQX, indicating the presence of AMPA/kainite receptors (**Fig 3K**).

### miR-124 and ISX9 exhibit both independent and cooperative transcriptional contributions in the reprogramming process to iNs

To in-depth analyze the molecular mechanism through which miR-124 contributes to the reprogramming process either alone or following ISX9 supplementation, we performed RNA-sequencing of miR-124-iNs and miR-124+ISX9-iNs at d7, using as controls astrocytes obtained the initial day of the reprogramming (d1) and sc-miRNA transfected astrocytes at d7. The differential expression analysis was performed between d7 miR-124-iNs or miR-124+ISX9-iNs and d1 astrocytes (astro) to uncover the whole spectrum of transcriptional changes occurring during neurogenic reprogramming of initiating astrocytes (miR-124-iNs *vs* astro and miR-124+ISX9-iNs *vs* astro respectively), whereas the d7 sc-miRNA-transfected astrocytes (sc-miRNA astro) were used the ultimate control for the identification of miR-124 target genes (see **Fig 5** below).

We identified 4,233 differentially expressed genes (DEGs) in miR-124-iNs *vs* astro and 6,652 DEGs in miR-124+ISX9-iNs *vs* astro (1≤log_2_(fold change)≤-1, FDR<0.05) (**Suppl Fig 4A**). Heat map analysis of DEGs (miR-124-iNs *vs* astro and miR-124+ISX9-iNs *vs* astro) belonging to the GO terms: Glial cell differentiation, Gliogenesis, Astrocyte development, Generation of neurons, Neuron differentiation, Regulation of neuron differentiation, Neurotransmitter transport and Synaptic signaling (**Fig 4A**) indicated that miR-124 alone efficiently down-regulated a gene cluster enriched in astrocytic genes (**Cluster I**). At the same time miR-124 up-regulated a gene cluster of neuronal specific genes (**Cluster III**) many of which were further up-regulated by ISX9 supplementation, while ISX9 highly up-regulated a neuronal specific gene cluster (**Cluster II**) that was most exclusively expressed in miR-124+ISX9-iNs.

**Figure 4:**
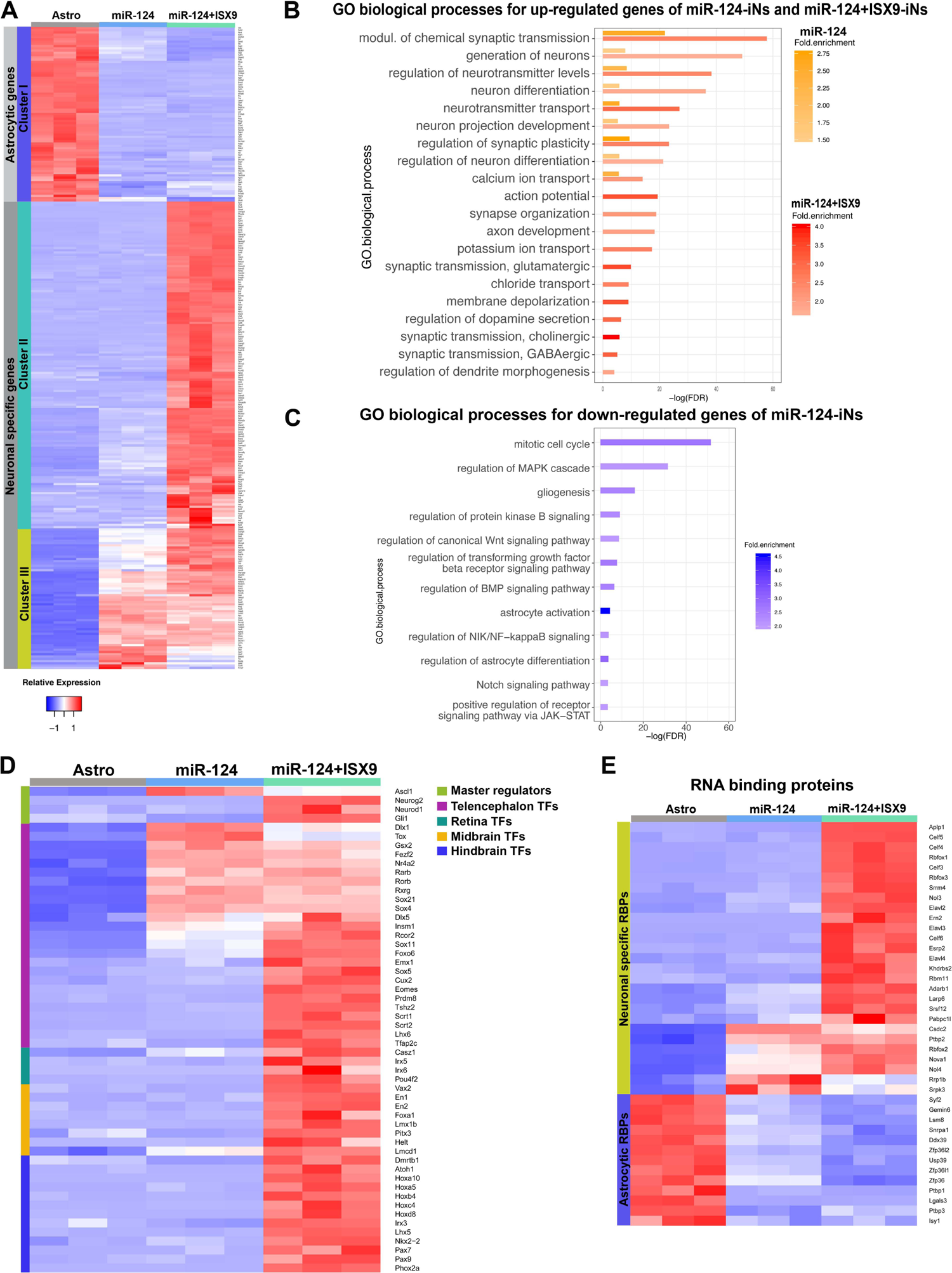
RNA-Seq analysis revealed both independent and cooperative transcriptional contributions of miR-124 and ISX9 in the reprogramming process. **(A)** Heat map analysis of 300 up- and down-regulated DEGs that belong to the GO terms: Glial cell differentiation, Gliogenesis, Astrocyte development, Generation of neurons, Neuron differentiation, Regulation of neuron differentiation, Neurotransmitter transport and Synaptic signaling. **(B)** GO analysis of biological processes for the up-regulated DEGs in miR-124-iNs *vs* astro (in orange) and miR-124+ISX9-iNs *vs* astro (in red). **(C)** GO analysis of biological processes for the down-regulated DEGs in miR-124-iNs *vs* astro. GO terms are ranked according to log_10_FDR and the intensity legends indicate the fold enrichment of each GO term. **(D)** Heat map analysis of 54 up-regulated differentially expressed TFs clustered according to the brain region they are developmentally expressed (telencephalon, retina, midbrain and hindbrain). **(E)** Heat map analysis of 40 up- and down-regulated RBPs.

GO enrichment analysis of biological processes for the up-regulated DEGs of both miR-124-iNs *vs* astro and miR-124+ISX9-iNs *vs* astro, further revealed that miR-124 alone up-regulated genes related to generation of neurons and neuronal differentiation as well as to more specific neuronal functions mostly associated to synaptic transmission (**Fig 4B in orange**), while ISX9 greatly enhanced the number of up-regulated genes related to more mature neuronal functions such as action potential, axon development and subtype specific synaptic transmission (**Fig 4B in red**). Furthermore, enrichment analysis of the down-regulated DEGs of miR-124-iNs *vs* astro indicated that many of them were related to cell cycle, gliogenesis, and astrocyte differentiation (**Fig 4C**). Interestingly, this analysis revealed a strong effect of miR-124 in down-regulating components of many signaling pathways, including MAPK, PKB, canonical Wnt, TGF-β, BMP, Notch and JAK/Stat signaling pathways (**Fig 4C**), known to play important roles in astrocytic identity and function (Gross et al. 1996; Acaz-Fonseca et al. 2019; Kang and Hébert 2011; Yang et al. 2012).

Since reprogramming is a process that implicates great changes in the transcriptomic, post-transcriptomic and epigenetic landscape of trans-differentiating cells, we sought to identify the differentially expressed transcription factors (TFs), epigenetic factors (EFs) and RNA binding proteins (RBPs) in our datasets that could possibly drive the reprogramming process. Heat map analysis of astrocytic TFs (**Suppl Fig 4Β**) indicated that miR-124 potently down-regulated TFs related to astrocytic function such as *Id1, Id3, Tcf4, Tcf7l1, Rbpj* and *Nfic,* while the addition of ISX9 exhibited minor contribution to their down-regulation (**Suppl Fig 4Β**). Importantly, validation of many of those genes with RT-qPCR verified the observed trend and also indicated that ISX9 alone (sc-miRNA+ISX9) failed to down-regulate their mRNA levels (**Suppl Fig 4C**). In parallel, heat map analysis of up-regulated neuronal-specific TFs revealed that miR-124 up-regulated TFs related to telencephalon development such as *Tox, Foxo6, Scrt1, Scrt2* and *Rcor2* (**Fig 4D**), along with TFs that we had already identified through prior qRT-PCR analysis (*Mash1, Insm1, Hes6, Sox4, Fezf2, Gsx2, Dlx1*). Additionally, ISX9 increased the number of TFs implicated in telencephalon development, among which *Eomes (Tbr2), Prdm8, Ovol2, Tfap2c* and *Tshz2* (**Fig 4D**), but unexpectedly it also highly up-regulated TFs related to more ventral/caudal brain regions, such as the retina, midbrain, hindbrain/spinal cord (**Fig 4D**), possibly implying that ISX9 expands region-specific neuronal identity at the transcriptional level. Validation of selected TFs/EFs expressed either in telencephalon (**Suppl Fig 4D**) or midbrain (**Suppl Fig 4E**) and hindbrain/spinal cord (**Suppl Fig 4F**) by RT-qPCR verified their up-regulation by ISX9.

Interestingly, heat map analysis of differentially expressed RNA-binding proteins (RBPs) revealed that miR-124 was sufficient to down-regulate many RBPs expressed in astrocytes such as the splicing factors *Ptbp1, Snrpa1, Lgals3* and *Isy1* and the mRNA decay proteins *Zfp36, Zfp36l1, Zfp36l2* (**Fig 4E**). In addition, miR-124 moderately up-regulated neuronal specific RBPs, among which *Elavl2, Elavl4, Nova1, Rbfox1, Rbfox2, Celf3* and *Nol3,* while ISX9 induced their further up-regulation and significantly increased the number of neuron specific RBPs such as *Aplp1, Celf4, Celf5, Celf6, Elavl3, Ern2, Esrp2* and *Rbfox3* (**Fig 4E**).

The analysis of differentially expressed EFs revealed that miR-124 increased the levels of several EFs related to epigenetic transcriptional activation, including *Kmt2c*, *Tet1*, *Smarca1, Smarca2, Chd7* and *Ss18l1 (Crest)* (**Suppl Fig 4G**). On the other hand ISX9 further contributed in up-regulating *Chd5, Actl6b*, *Dpf3* and *Smyd3*, while interestingly it majorly contributed in down-regulating EFs related to epigenetic transcriptional repression, such as *Suv39h1*, *Suv39h2*, *Hdac5* and *Hdac7* (**Suppl Fig 4G**).

The above observations led us conclude that miR-124 is sufficient to induce the astrocytic reprogramming switch towards an immature cortical neuronal fate through down-regulation of many glial-specific genes, many of which exhibit important regulatory functions. ISX9, on the other hand, acts auxiliary by enhancing neuronal-specific gene transcription contributing to the maturation of miR-124 immature-iNs.

### The non-neuronal RBP Zfp36l1 is a novel direct target of miR-124

To get a closer insight into the post-transcriptional effects of miR-124 on the astrocytic transcriptome, we sought to determine the direct targets of miR-124 that could act as drivers of the reprogramming switch. For this, we utilized publicly available AGO-HITS-CLIP data performed in mouse brain cortex (Chi et al. 2009) in order to detect miR-124 binding sites. The analysis revealed 171 miR-124 direct targets that were also defined as down-regulated in the miR-124-iNs *vs* sc-miRNA astro RNA-Seq analysis (log_2_(fold change)≤-1, FDR<0.01). miR-124 targets were subsequently filtered to examine genes expressed in astrocytes, utilizing a published reference list for expressed transcripts in distinct cell types of the mouse brain (Y. Zhang et al. 2014), ending up with 130 miR-124 direct target genes (**Fig 5A**). Interestingly, among these genes we identified the known targets of miR-124, Sox9 and Lhx2, but failed to detect Ptbp1 and Scp1, which are well characterized miR-124 targets with prominent role in neurogenic reprogramming.

**Figure 5:**
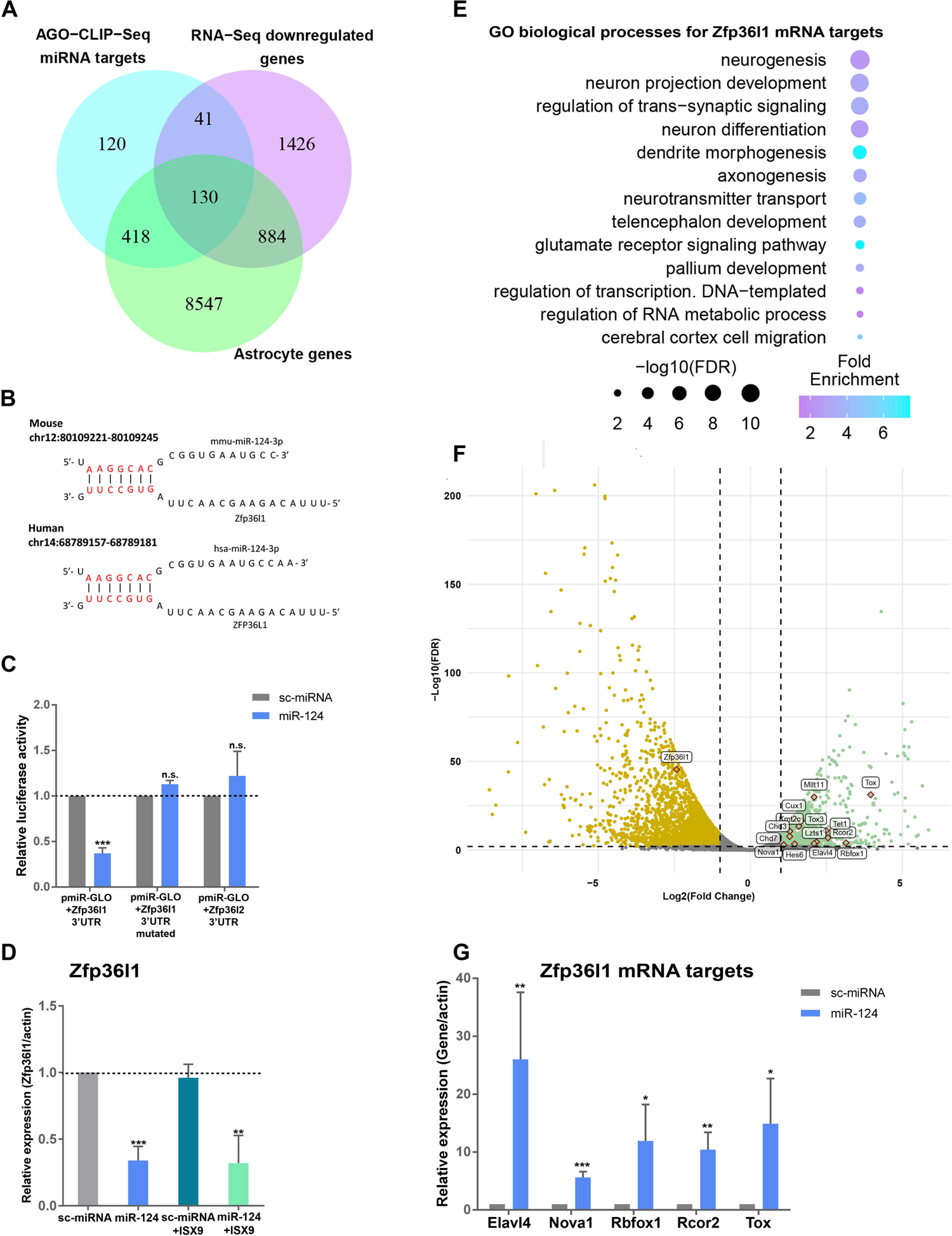
The RNA-binding protein Zfp36l1 is a novel direct target of miR-124 (Α) Venn diagram of the miR-124 targets, derived from AGO-HITS-CLIP and the down-regulated DEGs of miR-124 *vs* sc-miRNA astro RNA-Seq data (log2(fold change)≤-1, FDR<0.01). Identified miR-124 targets were combined with a public reference list of genes, expressed in astrocytes, resulting in a set of 130 genes. **(Β)** miR-124 direct binding to the 3’ UTR of Zfp36l1 and ZFP36L1, in mouse and human species respectively. miR-124 binds with perfect seed complementarity (7mer-m8 site) in both species. **(C)** Analysis of luciferase activity in HEK293T cells upon co-transfection with sc-miRNA or miR-124 mimics along with the reporter construct pmiR-GLO containing the 3’UTR of Zfp36l1 with the miR-124 binding site (n=7 independent experiments) or the 3’UTR of Zfp36l1 with the mutated miR-124 binding site (n=3) or the 3’UTR of Zfp36l2 (n=6). Normalized Firefly luciferase activity for sc-miRNA has been set to 1, (average ± SD). **(D)** RT-qPCR validation of the mRNA levels of miR-124 direct target *Zfp36l1* at d7 (average ±SD, n=3 independent experiments). **(E)** GO analysis of biological processes for Zfp36l1 direct targets that are also significantly up-regulated in miR-124-iNs *vs* sc-miRNA astro. GO terms are ranked according to log_10_FDR and the intensity legend shows the fold enrichment of each GO term. **(F)** Volcano plot comparing the log_2_(fold change) of TPM values in the miR-124-iNs vs sc-miRNA astro condition versus the log_10_(FDR) values. Significantly up-regulated (log_2_(fold change)≥1, FDR<0.05) and down-regulated (log_2_(fold change)≤-1, FDR<0.05) genes are shown in green and orange respectively. Labels of Zfp36l1 and neuronal-specific up-regulated genes that are also Zfp36l1 direct targets are portrayed. **(G)** RT-qPCR validation of the mRNA levels of the Zfp36l1 targets *Elavl4, Nova1, Rbofox1, Rcor2* and *Tox* at d7 (average ±SD, n=3 independent experiments). For all data presented *p<0.05, **p<0.01 and ***p<0.001 *vs* sc-miRNA.

In search of novel miR-124 targets with regulatory role, a prominent target was the RBP Zfp36l1, implicated in the mRNA decay (Lai et al. 2000) and highly expressed in cortical glial cells (Weng et al. 2019), cortical radial glial precursors (Yuzwa et al. 2017; Weng et al. 2019) and other non-neuronal cells (Carrick and Blackshear 2007). miR-124 directly binds to the 3’ UTR of Zfp36l1 transcript with perfect seed complementarity (7mer-m8 site) (**Fig 5B**), while in re-analysis of publicly available AGO-HITS-CLIP data from human motor cortex and cingulate gyrus tissues, the miR-124 binding site on the 3’ UTR of ZFP36L1 human transcript was found to be conserved (**Fig 5B**).

The targeting of this binding site in Zfp36l1 3’UTR by miR-124 was further validated by luciferase assays, where it was shown that its mutation completely blocked the strong down-regulation of luciferase activity induced by miR-124 (**Fig 5C, Suppl Fig 5A and B**). Of note, miR-124 does not target the 3’UTR of Zfp36l2, predicted to contain 2 binding sites (**Fig 5C and Suppl Fig 5C and D**), further supporting its specificity on Zfp36l1 regulation. The efficient down-regulation of *Zfp36l1* by miR-124 was also validated by RT-qPCR, revealing that it is a very early event in the course of reprogramming (**Suppl Fig 5E**). Interestingly, ISX9 was not potent to down-regulate *Zfp36l1* mRNA levels (**Fig 5D**), further supporting our initial observation that ISX9 alone cannot instruct reprogramming of astrocytes to iNs, possibly by failing to down-regulate astrocytic genes.

Since Zfp36l1 acts by mediating degradation of its mRNA targets, we were interested in identifying Zfp36l1 mRNA targets, being up-regulated in our analysis. For this purpose, we combined two publicly available Zfp36l1 iCLIP-Seq data from thymocytes (Vogel et al. 2016) and B lymphocytes (Galloway et al. 2016) and ended up with 621 Zfp36l1 direct targets that are up-regulated in miR-124-iNs *vs* sc-miRNA astro (log_2_(fold change)≥1, FDR<0.05), which importantly correspond to 47% of miR-124 up-regulated DEGs. GO enrichment analysis revealed that many of these genes are implicated in neurogenesis, neuron projection development, synaptic transmission, axonogenesis, dendritic morphogenesis and telencephalon development (**Fig 5E**). Interestingly, many of them were also found to regulate transcription and RNA processing (**Fig 5E**), highlighting an important regulatory role for many Zfp36l1 targets with possible impact on the reprogramming process. Among these targets we found neurogenic TFs, such as *Tox, Tox3, Rcor2, Cux1, Hes6, Lzts1* and *Mllt11*and EFs related to neurogenesis such as *Chd3, Chd7, Kmt2c* and *Tet1* (**Fig 5F**). Notably, we also identified as Zfp36l1 targets the neuronal RBPs *Elavl4, Nova1* and *Rbfox1* (**Fig 5F**). This constitutes a significant finding that suggests the repression of neuronal RBPs’ directly by Zfp36l1, being relieved upon miR-124-mediated Zfp36l1 down-regulation.

We subsequently examined the mRNA levels of the Zfp36l1 targets identified by our analysis, *Elavl*4, *Nova1, Rbfox1, Rcor2* and *Tox* upon miR-124 overexpression with RT-qPCR and verified that miR-124 was potent to induce their up-regulation (**Fig 5G**). We also verified the increase of the protein levels of the neurogenic TF, Tox, in miR-124-iNs at d5 (**Suppl Fig 5F and G**). Notably, we observed that the combination of miR-124 with ISX9 greatly further enhanced the mRNA levels of the RBP identified as targets of Zfp36l1, *Elavl4* and *Rbfox1* (**Suppl Fig 5H**), as well as the other neuronal RBPs of the nElavl family, *Elavl2* (**Suppl Fig 5I**) and *Elavl3* (**Suppl Fig 5J**), highlighting the importance of ISX9 in reinforcing and supplementing the neurogenic action of miR-124.

Collectively, these data show that the identified here targets of Zfp36l1 correspond to half of the miR-124-up-regulated genes and many of them constitute neuronal-specific genes with important regulatory functions. Thus the miR-124/Zfp36l1 interaction bears the potential to play a key regulatory role in the control of the neurogenic reprogramming switch of astrocytes by miR-124.

### Targeting of Zfp36l1 by miR-124 plays a key role in the miR-124-induced cell fate switch of astrocytes to iNs

To investigate the impact of the miR-124/Zfp36l1 interaction on the miR-124-mediated reprogramming of astrocytes, we used a target site blocker (TSB) oligonucleotide that competitively binds to miR-124 binding site on the 3’ UTR of Zfp36l1 mRNA, blocking its down-regulation by the miR-124-RISC complex (**Fig 6A**). The moderate down-regulation of Zfp36l1 protein by miR-124 as early as d5 of reprogramming was reversed by TSB at the miR-124:TSB molecular ratio 2:1, as estimated by western blot (**Fig 6B and C**), while we observed no significant effect of TSB on Zfp36l2 protein levels (**Suppl Fig 6A and B**). Due to extensive stress and cell death, the effect of TSB on the reversal of Zfp36l1 protein levels upon miR-124 over-expression was not possible to be estimated for a longer time point of the reprogramming.

**Figure 6:**
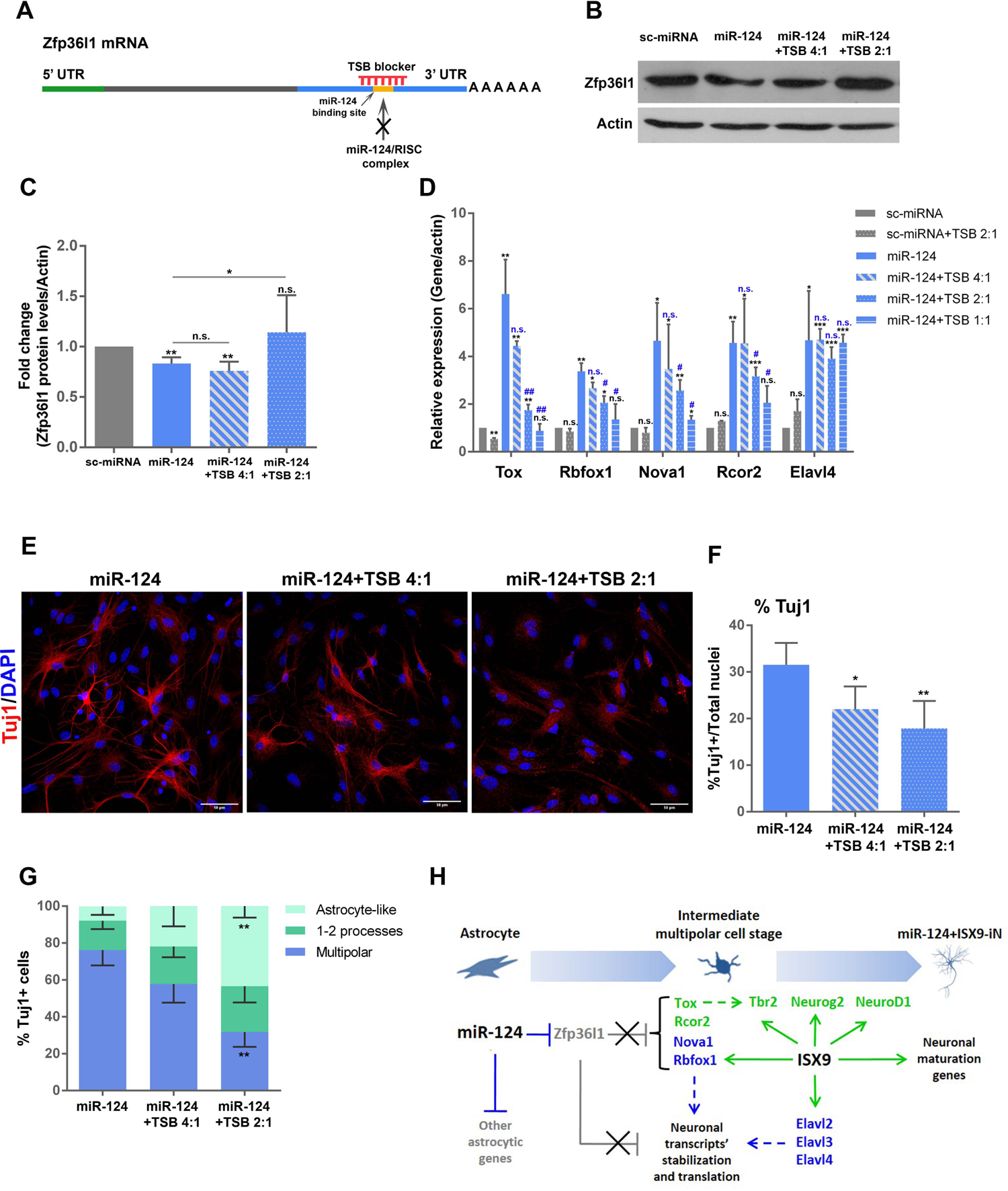
Targeting of Zfp36l1 by miR-124 plays a key role in the miR-124-induced cell fate switch of astrocytes to iNs. **(A)** Schematic representation of the TSB binding region in the 3’UTR of *Zfp36l1* mRNA. **(B)** Western blot analysis of the protein levels of Zfp36l1 in the absence or presence of increasing concentrations of TSB (miR-124:TSB molecular ratio 4:1 and 2:1) at d5 of the reprogramming protocol. Actin protein levels have been used as loading control. **(C)** Quantification of Zfp36l1 protein levels normalized to β-actin. Normalized Zfp36l1 protein levels in sc-miRNA transfected astrocytes have been set to 1, (average ± SD, n=3 independent experiments, *p<0.05, **p<0.01 *vs* sc-miRNA). **(D)** RT-qPCR analysis of the mRNA levels of the Zfp36l1 targets *Tox, Rbfox1, Nova1, Rcor2* and *Elavl4* in the presence or absence of increasing concentrations of TSB (miR-124:TSB molecular ratio 4:1, 2:1 and 1:1) at d3 of the reprogramming protocol (average ± SD, n=3 independent experiments,*p<0.05, **p<0.01 and ***p<0.001 *vs* sc-miRNA and ^#^p<0.05, ^##^p<0.01 *vs* miR-124). **(E)** Immunostaining of astrocytes reprogrammed with miR-124 or miR-124 along with increasing concentrations of TSB (4:1 and 2:1) with anti-Tuj1 antibody at d5 of the reprogramming protocol. **(F)** Quantification of the percentage of Tuj1+ cells at d5 (average ±SD, n=3 independent experiments, *p<0.05 and **p<0.01 *vs* miR-124). **(G)** Morphological characterization of Tuj1+ cells at d5. Quantification of the percentages of multipolar Tuj1+ cells (3 or more processes extending from soma), Tuj1+ cells with 1-2 processes and Tuj1+ cells with an astrocyte-like morphology (rectangular soma with none or 1-2 processes) (average ± SD, n=3 independent experiments, **p<0.01 *vs* miR-124). **(H)** Proposed model of independent and cooperative transcriptional and post-transcriptional contributions of miR-124 and ISX9 in miR-124/ISX9– induced direct neuronal conversion of cortical astrocytes. Post-transcriptional events and the RBPs are highlighted in blue, transcriptional events along with the TFs are shown in green, while down-regulated genes and (blocked) inhibitory mechanisms are shown in grey, (dashed lines present knowledge from the literature).

Next, we tested the effect of TSB on the mRNA levels of the identified neuronal targets of Zfp36l1, *Tox, Rbfox1, Nova1, Rcor2* and *Elavl4* by RT-qPCR initially at d3 and observed a dose-dependent reduction of the miR-124-induced up-regulation of *Tox, Rbfox1, Nova1* and *Rcor2*, but not of *Elavl4*, in the presence of increasing concentrations of TSB (miR-124:TSB molecular ratio 4:1, 2:1 and 1:1) (**Fig 6D**). The mRNA levels of *Tox* and *Rbfox1* retained the same response to TSB at a later time point, at d5 (**Suppl Fig 6C and D**), an effect that was not observed for *Nova1*, *Rcor2* and *Elavl4* (**Suppl Fig 6E**), implying that their post-transcriptional regulation becomes more complex as the neuronal conversion is gradually established. Importantly, TSB resulted in a significant and dose-dependent reduction in the percentage of Tuj1+ miR-124-iNs at d5 (**Fig 6E and F**), with an evident alteration of their characteristic immature neuronal morphology (**Fig 6E and G**). More specifically, morphological analysis of the number of processes extending from the soma and the size of the soma of Tuj1+ cells indicated that TSB addition resulted in gradual abolishment of their multipolar morphology with fine processes, characteristic of miR-124-iNs (**Fig 6E left panel and G**) and instead largely led to the retention of a premature astrocyte-like morphology with bigger soma (**Suppl Fig 6F**) and none or very few processes (**Fig 6E right panel and G**).

These observations strongly indicate that the targeting of Zfp36l1 by miR-124 is an important event during the miR-124-induced reprogramming by unlocking neurogenic genes with important regulatory activity and by controlling the necessary morphological changes.

### miR-124 induces reprogramming of resident reactive astrocytes to immature iNs *in vivo* following cortical trauma

To evaluate the *in vivo* reprogramming potential of miR-124 to drive cell fate conversion of resident reactive cortical astrocytes to iNs alone or in synergy with ISX9, we applied two viral-based approaches (using lentiviral and AAV vectors) to overexpress miR-124 into the mechanically injured cortex of 3-4 months old mice. In the lentiviral transfer approach, we stereotaxically injected either control LV-GFP or LV-miR-124-GFP to transduce reactive astrocytes surrounding the cortical injury site, while ISX9 was systemically administered for 5 consecutive days beginning 2 days after viral injection (p.i.) in a group of mice (**Fig 7A**). Seven days p.i. the majority of transduced cells surrounding the injury site (**Suppl Fig 7A**) in both conditions were GFAP+ reactive astrocytes (71.6% ± 8.0% for LV-GFP and 65.7% ± 15.7% for LV-miR-124-GFP), while the percentage of transduced NeuN+ neurons was much lower (6.4% ± 3.6% for LV-GFP and 15.3% ± 3.7% for LV-miR-124-GFP) (**Suppl Fig 7B**). Analysis of NeuN+/GFP+ transduced cells 3weeks (3w) p.i. revealed that 71.8% ± 12.4% of the LV-miR-124-transduced cells were NeuN+, while only 10.2% ± 2.6% of the LV-GFP transduced cells were NeuN+ exhibiting no significant difference between the 7d and 3w time points (**Fig 7B and C**). These results point to a strong potential of miR-124 to drive direct conversion of reactive astrocytes into iNs *in vivo*, while the administration of ISX9 showed no further significant contribution to this conversion (**Fig 7C**). The vast majority of NeuN+ iNs in both groups were also positive for the cortical marker Tbr1 (98% ± 4.1% for LV-miR-124-GFP and 88% ± 11.1% for LV-miR-124-GFP+ISX9) (**Suppl Fig 7C and D**).

**Figure 7:**
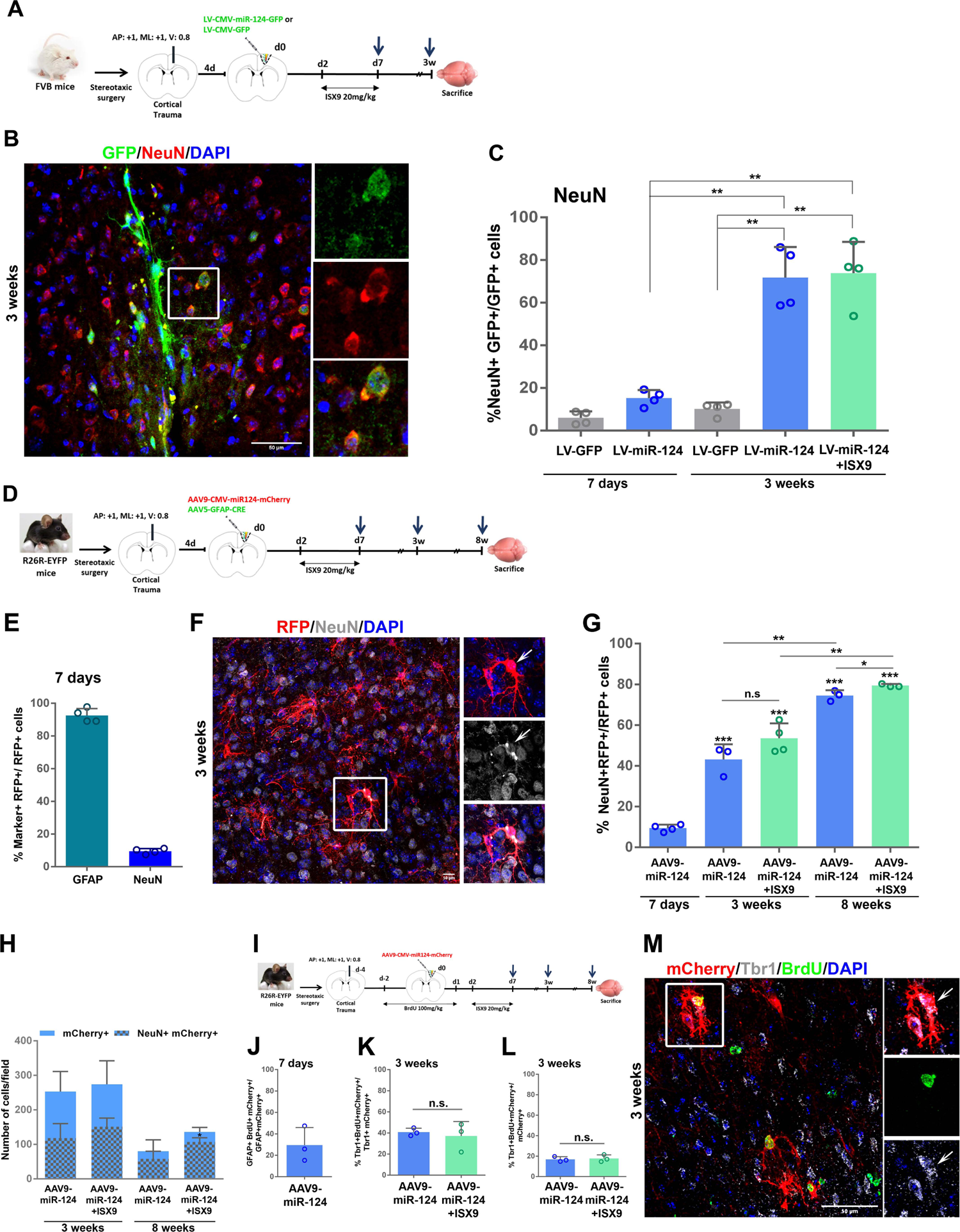
miR-124 induces reprogramming of resident reactive astrocytes to iNs with cortical identity *in vivo* following cortical trauma. **(A)** Experimental setup for the lentiviral approach. **(B)** LV-124-transduced cells in the peritraumatic cortical parenchyma expressing the mature neuronal marker NeuN, 3w after viral transduction. Inset area indicated in white frame. **(C)** Percentage of control LV-GFP and LV-124-transduced cells expressing NeuN, 7d and 3w post-transduction with or without treatment with ISX9, (average ± SD, n=4 animals for all groups and time points, **p<0.01). **(D)** Experimental setup for the AAV approach. **(E)** Percentage of AAV9-miR-124-transduced cells expressing the astrocytic marker GFAP or the mature neuronal marker NeuN, 7d p.i. (average ± SD, n=3 animals). **(F)** AAV9-124-transduced cells (in red detected with an anti-RFP antibody) expressing NeuN, 3w p.i. Inset area indicated in white frame. **(G)** Percentage of AAV9-miR-124-transduced cells in the peritraumatic parenchyma expressing NeuN at d7, 3w and 8wp.i. with or without treatment with ISX9, (average ± SD, n=3-4 animals per group, *p<0.05, **p<0.01 and ***p<0.001). **(Η)** Representation of the average number of mCherry+ (blue bars) and NeuN+/mCherry+ cells (inner gray-blue bars) present in the peritraumatic parenchyma 3w and 8w p.i. in the presence or absence of ISX9, (average ± SD, n=3 animals per group per time point, *p<0.05 for the number of NeuN+/mCherry+ cells in AAV9-miR-124+ISX9 *vs* AAV9-miR-124 at 8w). **(I)** Experimental setup for the BrdU administration. **(J)** Percentage of GFAP+/mCherry+ cells that have incorporated BrdU 7d p.i. (average ± SD, n=3 animals). Percentage of Tbr1+/mCherry+ cells that originated from a proliferating state (BrdU+) **(K)** and percentage of mCherry+ cells that originated from a proliferating state and have undergone reprogramming (Tbr1+/BrdU+) **(L)** 3w p.i. (average ± SD, n=3 animals per group). **(M)** miR-124-transduced cells (in red detected with an anti-mCherry antibody) in the peritraumatic parenchyma that express Tbr1 and have incorporated BrdU 3w p.i. Inset area indicated in white frame.

To further validate the astrocytic origin of iNs upon miR-124 force-expression in the same cortical trauma model, supplementary to the lentiviral approach we employed a viral lineage tracing strategy by stereotaxically co-injecting AAV5-GFAP-Cre virus, to mark cortical astrocytes, along with AAV9-CMV-miR-124-mCherry virus to overexpress miR-124 in the cells of the injured cortex of R26R-EYFP reporter mice (**Fig 7D**). However we observed unexpected GFP expression in peritraumatic resident neurons besides its expression in astrocytes as early as 7d after AAV5-GFAP-Cre injection, indicating that in accordance with recent reports (Wang et al. 2021), this approach cannot warranty exclusive astrocytic lineage tracing. Given this limitation, we restricted our analysis to the AAV9-CMV-miR-124-expressing cells, also employing BrdU administration to label the proliferating astrocytes’ population of the peritraumatic area. Analysis of the cell type of AAV9-miR-124-transduced cells 7d p.i. revealed that their majority was GFAP+ reactive astrocytes (92.5% ± 4.2%), with only 9.5% ± 2.1% being NeuN+ neurons (**Fig 7E and Suppl Fig 8A**), while 3w later 42.8% ± 7.8% of them were NeuN+ iNs, exhibiting an immature neuronal morphology (**Fig 7F and G, Suppl Fig 9B**). ISX9 administration along with miR-124 led to a small, not statistically significant increase in NeuN+ iNs’ percentage (51.6% ± 7.5%) (**Fig 7G**), without further affecting their morphology. In accordance with the lentiviral approach, in both conditions the percentage of Tbr1+ iNs was almost equal to that of NeuN+ iNs (41.2% ± 3.6% in AAV9-miR-124 and 51.4% ± 15.4% in AAV9-miR-124+ISX9) (**Suppl Fig 8B**). Further analysis at the longer time point of 8w revealed that the percentage of NeuN+ iNs increased to 74.5% ± 2.6% (**Fig 7G**), with a small but statistically significant additional increase (79.4% ± 0.8%) in AAV-miR-124+ISX9 (**Fig 7G**). Still the average number of remaining NeuN+ iNs per section was 50% lower in AAV9-miR-124 (58 ± 22 at 8w *vs* 118 ± 42 at 3w p.i.) and 30% lower in miR-124+ISX9 (107 ± 12 at 8w *vs* 151 ± 25 at 3wp.i.) **(Fig 7H)**. Interestingly the average number of NeuN+ iNs in the AA9-miR-124+ISX9 group was almost double as compared to the AAV9-miR-124 group (107 ± 12 for AAV9-miR-124+ISX9 *vs*58 ± 22 for AAV9-miR-124) **(Fig 7H),** indicating that ISX9 enhances the survival of converted iNs. However, the majority of reprogrammed iNs in both conditions retained an immature neuronal morphology even at the time point of 8wp.i. (**Suppl Fig 8C and Suppl Fig 9F and G**). Of note, mCherry expression was also observed in several resident pyramidal neurons mostly in the broader peritraumatic region at both time points, a phenomenon very recently pointed out for AAVs (Wang et al. 2021), however mCherry+ resident pyramidal neurons were excluded from our analysis as they were easily distinguished by their morphology (large conic soma and large apical process) (**Suppl Fig 8D and Suppl Fig 9C**).

Further on, to validate that produced iNs are the result of a reprogramming process, we administered BrdU for 4 consecutive days starting 2 days after the cortical trauma to label proliferating reactive astrocytes of the peritraumatic area (**Fig 7I**). Analysis 7 dp.i. revealed that nearly 1/3 of transduced astrocytes (29.5 ± 16.3%) had incorporated BrdU (**Fig 7J**), while BrdU+ transduced cells were still present 3w p.i. in both conditions (34.5% ± 5.5% for AAV9-miR-124 and 33.5% ± 12.7% for AAV9-miR-124+ISX9) (**Suppl Fig 8E**). Importantly, 3w p.i. iNs originating from a proliferating state expressing the cortical marker Tbr1were present (**Fig 7M and Suppl Fig 9D and E)** at a percentage of 40.8% ± 3.7% in AAV9-miR-124 and 37.2% ± 13.6% in AAV9-miR-124+ISX9 cells (**Fig 7K**) and accounted for nearly 1/5 of all transduced cells’ population (**Fig 7L**). Of note, BrdU+/Tbr1+ transduced cells were still present at 8w p.i. (**Suppl Fig 8F**).

These findings collectively indicate that a significant subpopulation of iNs originates from a proliferating fraction of reactive astrocytes. They also show that miR-124 is sufficient to drive on its own neurogenic reprogramming of reactive astrocytes to immature iNs which remain viable within the injured cortex for a period of at least 8w, while ISX9 administration seems to enhance the survival of converted iNs.

## Discussion

In this study we attempted to isolate miR-124 mechanism of action from that of other reprogramming co-factors and aimed at identifying novel miR-124 targets with important contribution in the reprogramming process. We provide evidence that miR-124 drives the trans-differentiation switch of cortical astrocytes to an immature iN phenotype of cortical identity, directing astrocytes through a multipolar intermediate stage expressing neurogenic TFs associated with neuronal commitment and differentiation during embryonic (Kwan, Šestan, and Anton 2012; Elsen et al. 2018) and adult neurogenesis (Díaz-Guerra et al. 2013). In order to overcome the unexpected finding that miR-124 over-expression significantly down-regulates the neuronal differentiation TF NeuroD1 (Z. Gao et al. 2009), we supplemented miR-124 with the neurogenic compound ISX9, known to up-regulate neuronal specific genes, among which NeuroD1. ISX9 acts by increasing intracellular Ca^2+^ signaling leading to HDAC5 nuclear exit and MEF2 de-repression (Schneider et al. 2008) and has been already used in chemical reprogramming protocols for the conversion of fibroblasts and astrocytes to iNs (L. Gao et al. 2017; Li et al. 2015). Indeed, the addition of ISX9 greatly improved both the reprogramming efficiency and differentiation status of miR-124+ISX9-iNs *in vitro*, leading to the acquisition of electrophysiologically active iNs. Interestingly, despite up-regulation of NeuroD1 levels and many other neuronal TFs by ISX9 alone, control sc-miRNA+ISX9-treated astrocytes failed to undergo reprogramming. NeuroD1 has been extensively reported to possess a strong ‘pioneer factor’ reprogramming capacity towards the neuronal fate upon its viral-mediated overexpression (Pataskar et al. 2016; Matsuda et al. 2019), however very high levels of NeuroD1–not triggered by ISX9 supplementation in our system– seem to be required to inflict these effects.

The RNA-Seq analysis we performed, highlighted the importance of miR-124 over-expression in the down-regulation of astrocytic TFs, RBPs, EFs and components of signaling pathways with significant regulatory role in astrocytic identity and function, which is in accordance with its documented role in favoring neuronal fate at the expense of astrocytic (Neo et al. 2014) and implies that astrocytic identity barriers need to be repressed before the induction of a neuronal cell fate during the reprogramming process. Conversely, ISX9 had a small contribution in the repression of astrocytic genes, thus this might be the reason for its inability to confer the reprogramming switch on its own.

Our analysis of the direct miR-124 targets utilizing AGO-HITS-CLIP data from mouse cortex(Chi et al. 2009) revealed the RBP *Zfp36l1* as a novel target of miR-124. Of note, we also verified the miR-124/Zfp36l1 interaction in human bearing the same binding site, by analyzing AGO-HITS-CLIP data from human motor cortex and cingulate gyrus (Boudreau et al. 2014), a finding highlighting the importance of this conserved interaction during mammalian brain evolution. Many studies have identified several direct targets of miR-124 with important regulatory role in neurogenesis, acting at the transcriptional level, such as the TFs *Sox9* (Cheng et al. 2009), *Lhx2* (Sanuki et al. 2011) and the components of the REST repressor complex, *Scp1* (Visvanathan et al. 2007) and *Rcor1* (Baudet et al. 2012; Volvert et al. 2014); or at the epigenetic level, such as the component of the PRC2 complex *Ezh2* (Neo et al. 2014) and the component of the BAF complex *BAF53a* (Yoo et al. 2009); as well as at the post-transcriptional level, such as the RBP involved in alternative splicing *Ptbp1*( Makeyev et al. 2007).

Here, we report for the first time that miR-124 is directly implicated in the regulation of another process mediated by RBPs, the mRNA decay, apart from the well-characterized miR-124/Ptbp1 circuitry (Yeom et al. 2018; Xue et al. 2013; Makeyev et al. 2007). Zfp36l1 is a member of the Zfp36 family of proteins along with Zfp36 and Zfp36l2, which act by binding to AU rich elements (AREs) in the 3’UTR of their mRNA targets mediating their destabilization (Lai et al. 2000). Zfp36l1 is expressed in cortical radial glial precursors of the VZ (DeBoer et al. 2013; Yuzwa et al. 2017; Weng et al. 2019), cortical glial cells (Weng et al. 2019) and non-neuronal cells (M. T. Chen et al. 2015; Carrick and Blackshear 2007). Interestingly, the ARE-dependent mRNA decay is regulated by other neurogenic miRNAs as well, since the close paralog of Zfp36l1, Zfp36, is targeted by miR-9 (Dai et al. 2015), suggesting a combined regulation of Zfp36 family members by the two miRNAs to increase the stabilization of neuronal mRNAs during neurogenesis.

In order to identify targets of Zfp36l1 being up-regulated in our system we examined data from Zfp36l1-iCLIP-Seq experiments in thymocytes (Vogel et al. 2016) and B lymphocytes (Galloway et al. 2016). Although our analysis was restricted to a non astrocytic transcriptome, we identified a rather large number of Zfp36l1 targets being up-regulated in miR-124-iNs that interestingly correspond to nearly half of the up-regulated genes by miR-124. Importantly, many Zfp36l1 targets exhibit significant regulatory role in neurogenesis and neuronal differentiation, such as the studied here TFs *Tox* and *Rcor2* and the RBPs *Rbfox1*, *Elavl4* and *Nova1*. Having verified that Zfp36l1, and not Zfp36l2, is a direct target of miR-124, we used a target site blocker (TSB) that efficiently antagonizes the binding of miR-124 in the 3’UTR of *Zfp36l1* mRNA, to investigate whether Zfp36l1 targeting by miR-124 contributes to the cell fate switch of astrocytes. Indeed the disruption of miR-124/Zfp36l1 interaction had a negative impact on the reprogramming process, reducing the number of Tuj1+ miR-124-iNs and abolishing their multipolar morphology. Of note, among the neuron-specific RBPs identified as down-stream targets of Zfp36l1, *Rbfox1* and *Nova1* exhibit a pattern of direct post-transcriptional regulation in contrast to *Elavl4*, implying the stronger involvement of other RBPs in its post-transcriptional regulation. In accordance with this, Zfp36 has been reported to target *Elavl2, Elavl3, Elavl4* and *Nova1* (Dai et al. 2015), uncovering a complementary and synergistic role of Zfp36l1 and Zfp36in repressing neuron-specific RBPs in non-neuronal cells.

Furthermore, the addition of ISX9 significantly contributed to the robust up-regulation of the mRNA levels of *Rbfox1*, *Elavl2, Elavl4* and most prominently Elavl3, reinforcing the action of miR-124 in inducing the switch from the neuronal transcripts’ destabilizing RBPs to the stabilizing neuronal RBPs (Ince-Dunn et al. 2012; Weyn-Vanhentenryck et al. 2014, Scheckel et al. 2016) through a different mechanism that would be interesting to be explored (**Fig 6H**). Interestingly, miR-124 has been recently shown to synergize with Elavl3 for the enhancement of target gene expression during neuronal maturation (Lu et al. 2021), further supporting our observations for the role of ISX9 in promoting iNs’ maturation in cooperation with miR-124. In parallel, ISX9 supplementation greatly enhanced the transcriptional levels and the number of TFs related to telencephalon development and/or adult neurogenesis, further promoting the cortical identity of iNs already initiated by miR-124. Surprisingly, ISX9 also up-regulated a large set of TFs related to the development of other non-telencephalic brain regions such as the midbrain and the hindbrain/spinal cord. Of note, ISX9 has been shown to affect the epigenetic landscape leading to an open chromatin state by increasing H3/H4 acetylation in pancreatic β-cells (Dioum et al. 2011), suggesting that an epigenetic mechanism may be responsible for the observed transcriptional regional expansion, a hypothesis that needs to be further explored.

Here we also explored the *in vivo* capacity of miR-124 to induce direct neurogenic reprogramming of astrocytes in a mouse cortical injury model, known to render astrocytes more plastic by activating NSC genes’ expression, thus facilitating their reprogramming (Götz et al. 2015). *In vivo* reprogramming of reactive astrocytes to neuronal precursors and iNs has been achieved following forced expression of TFs among which Sox2 (Niu et al. 2013), Neurog2 (Grande et al. 2013), NeuroD1 (Rivetti Di Val Cervo et al. 2017; Guo et al. 2014) and Nurr1/Neurog2 (Mattugini et al. 2019), in some cases combined with anti-apoptotic and/or anti-oxidant treatment to enhance survival (Gascón et al. 2016), however the *in vivo* neurogenic capacity of miR-124 on its own has not been evaluated up to now. For this, we employed two different viral-based approaches, using either lentiviruses or the more recently used AAVs and corroborated the latter approach with BrdU administration during the peak period of trauma-induced astrogliosis. The last years, the functional restoration of disease-related symptoms upon direct reprogramming has been reported (Giehrl-Schwab et al. 2022; Qian et al. 2020; Zhou et al. 2020; Z. Wu et al. 2020; Y. C. Chen et al. 2019), supporting the therapeutic promise of *in vivo* direct reprogramming to restore impaired neuronal circuits (Bocchi, Masserdotti, and Götz 2022). However, certain AAV-based strategies and astrocyte-lineage tracing approaches have been very recently put under question as to their capacity to confer neurogenic reprogramming (Wang et al. 2021). In light of this new challenge in the field of *in vivo* reprogramming, we report here that both viral approaches point towards a reprogramming capacity of miR-124 in converting reactive astrocytes of the peritraumatic cortical area to iNs of cortical identity. More importantly, our AAV-based protocol combined with BrdU administration, which is one of the very few widely accepted tools currently available to verify the non-neuronal origin of reprogrammed cells, showed that about 1/3 of iNs originated from proliferating astrocytes, strongly supporting the reprogramming potential of miR-124 towards NeuN+/Tbr1+ iNs – harboring however an immature phenotype – that persist in the peritraumatic area 8w following miR-124 over-expression. Additionally, our analysis revealed that ISX9 supplementation seems to confer a survival advantage to the converted iNs, implying a cell- or non-cell autonomous contribution of ISX9 in iNs’ survival that needs further exploration.

These findings indicate that, similarly to the *in vitro* situation, miR-124 is capable of inducing the reprogramming switch of peritraumatic reactive astrocytes, but is not sufficient to further enhance the maturation of produced immature iNs. Additionally, ISX9 *in vivo* administration acts beneficially to reprogramming, by contributing majorly to iNs’ enhanced survival for a longer period, but in contrast to our *in vitro* observations it does not seem to reinforce iNs’ maturation, although that it is well accepted that it crosses the blood brain barrier in the scheme that we administered it (Bettio et al. 2016). Further analysis of miR-124-iNs’ and miR-124+ISX9-iNs’ molecular profiling using single cell RNA-Seq could shed light on the exact differentiation state of miR-124-iNs and the possible enhancement of miR-124+ISX9-iNs’ maturation status by ISX9.

### Ideas and speculation

Taken together this study highlights the strong potency of miR-124 to instruct the cell fate switch of astrocytes, post-transcriptionally triggering cortical neurogenesis pathways being unlocked in part by the direct targeting of *Zfp36l1*. Additionally, our *in vitro* results give mechanistic insight into the combined action of miR-124 and ISX9 in driving direct reprogramming of astrocytes to mature iNs. Importantly, our findings point to the *in vivo* capacity of miR-124 to induce reactive astrocytes’ reprogramming to immature iNs able to survive for long periods, in particular following ISX9 supplementation, further supporting miR-124 ‘ master reprogramming’ potential. However, the synergistic effect of miR-124/ISX9 does not seem to be sufficient to drive iNs’ full maturation, indicating that more intrinsic and/or extrinsic cues are required. To this end the findings of our *in vitro* mechanistic studies point to certain common transcriptional and post-transcriptional downstream effectors of miR-124 and ISX9 that are known to promote neurogenesis and neuronal differentiation and thus bear the potential to amplify miR-124/ISX9 combined *in vivo* reprogramming capacity and to enhance the differentiation/ maturation of newly produced iNs. This strategy comes in accordance with a very recent view in the reprogramming field arguing that *in vivo* glia-to-neuron conversion is a two-stage process involving initial immature iNs production and their subsequent differentiation provided that certain still largely unknown barriers are over-passed (Leaman, Marichal, and Berninger 2022). Thus focusing on the points of convergence of miR-124/ISX9 action holds promise for the establishment of a more efficient, *in vivo* reprogramming protocol to improve maturation of iNs and their functional integration in the host circuit following trauma or neurodegeneration.

## Materials and Methods

### Primary cultures of postnatal cortical astrocytes

Primary postnatal astrocytic cultures from P3-P5 mice were prepared as previously described(Aravantinou-Fatorou et al. 2015). Briefly, the cerebral cortexes from 2-3 P3-P5 C57BL/6 mice were collected in ice cold HBSS (Invitrogen), the tissue was washed three times with HBSS and digested with 0.04% trypsin (Sigma) and 10μg/ml DNAse (Sigma) for 5 min at 37°C. After digestion cells were mechanically dissociated, centrifuged for 5 min at 850 rpm (120 g), re-suspended in DMEM 4.5g/lt glucose (Invitrogen) containing 10% FBS (Invitrogen), 1% Penicillin/Streptomycin (Pen/Strep) (Sigma) and placed in a T75 flask pre-coated with poly-D-lysine (PDL) (Sigma). When culture reached confluence (usually after 7 days), the flask was shaken in a horizontal shaker at 200-250 rpm for 20h, in order to obtain a pure astrocytic culture, free from neurons, oligodendrocytes and microglia. The remaining cells were digested with 0.5% trypsin-EDTA (Invitrogen) for 5 min at 37°C, centrifuged at 850 rpm, re-suspended in fresh DMEM 4.5g/lt glucose 10% FBS, 1% Pen/Strep and divided in two new T75 flasks pre-coated with PDL. Half of the medium was changed every two days.

### In vitro reprogramming protocol

For the reprogramming of astrocytes to induced-neurons, 40,000 astrocytes were seeded in 10mm coverslips coated with 20μg/ml poly-L-ornithine (PLO) (Sigma) overnight and 5μg/ml laminin for 3h at 37°C (Sigma). Once cells reached>90% confluence (usually after 1-2 days) they became transfected with 80nM miR-124-3p mimics or scrambled (sc-miRNA) mimics (negative control) (Thermo) using Lipofectamine 2000 (Invitrogen) according to manufacturer’s instructions (d1). The next day the astrocytic medium (DMEM 4.5g/lt glucose, 10% FBS, 1% Pen/Strep) was replaced with the reprogramming medium: Neurobasal (Invitrogen) supplemented with 1X B-27 (Invitrogen), 1X GlutaMAX (Invitrogen), 20μΜ vitamin E (a-tocopherol) (Sigma) and 200mM ascorbic acid (Sigma). The same process of transfection was repeated twice at d3 and d5. Vitamin E was added to the medium until d4, while ascorbic acid was added throughout the reprogramming protocol. At d7 the reprogramming medium was changed to the neuronal differentiation medium: Neurobasal supplemented with 1X B-27, 1X GlutaMAX, 20ng/ml BDNF (R&D Systems), 0.5mM cAMP (Sigma) and 200mM ascorbic acid. In the miR-124+ISX9-reprogrammed cells, 10μΜ of ISX9 chemical compound (Tocris) were added from d2 to d10. All the mediums added to the reprogrammed cells were pre-conditioned for 24h in a confluent astrocytic culture.

For BrdU administration, astrocytes were treated with 10μΜ BrdU (Sigma) two times per day from day1 until day4 of the reprogramming protocol and analyzed at d7.

### RT-qPCR analysis

For the RT-qPCR analysis experiments, total RNA was extracted using the Nucleospin miRNA kit (Macherey-Nagel) and 500-800ng of total RNA were used for cDNA synthesis with the Superscript II reverse transcriptase (Invitrogen) according to manufacturer’s instructions. Quantitative real time PCR was performed using SYBR Select Master Mix (Applied Biosystems) and samples were run in the ViiA 7 Real-Time PCR System (Applied Biosystems). The primers used are listed in **Table 1**. Each sample was analyzed in triplicates, gene expression was calculated using the ΔΔCt method and all the results were normalized to β-actin expression. Relative expression was estimated setting the values of sc-miRNA transfected astrocytes to 1. All experiments were performed at least in triplicates.

**Table 1:**
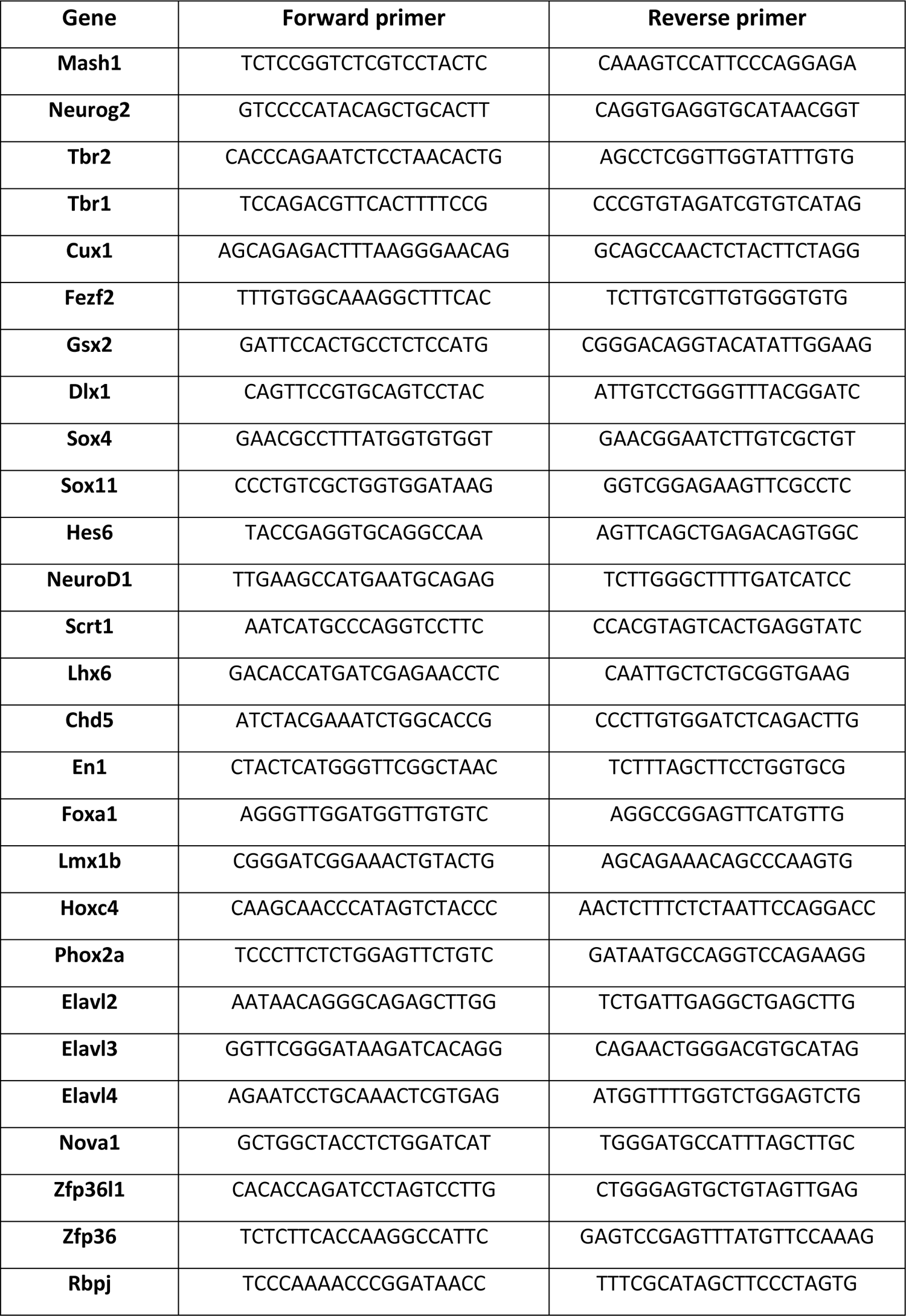
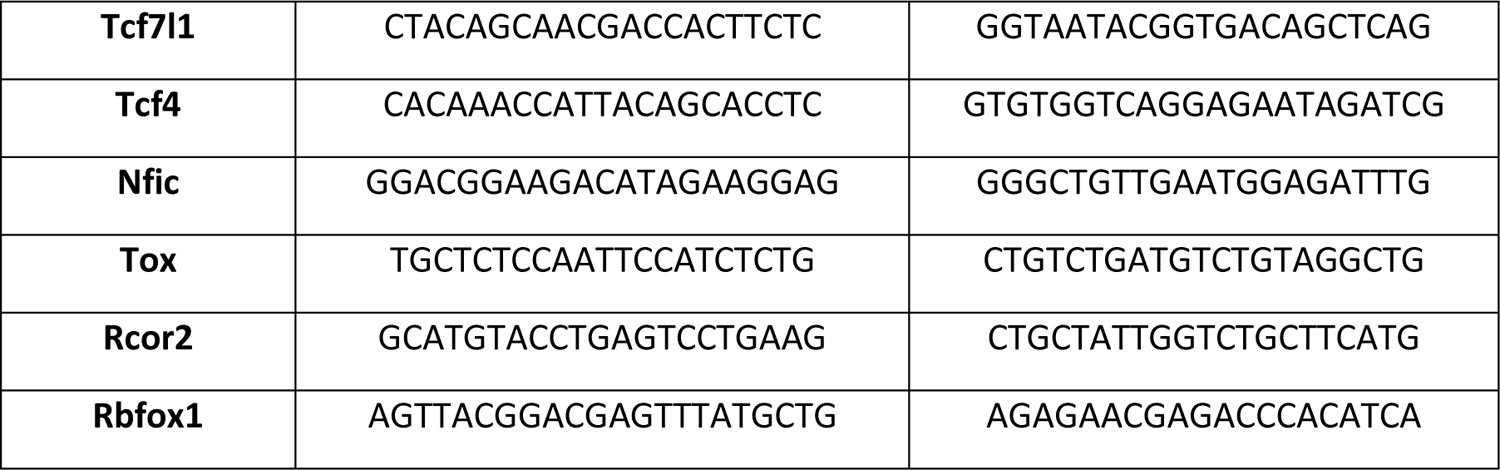
Sequences of primers for real-time PCR used in this study

### Immunocytochemistry

Cells were washed once with PBS and then fixed with 4% paraformaldehyde for 20 min at room temperature. Afterwards, cells were washed three times with PBS and blocked with 5% normal donkey serum (NDS) (Merck-Millipore), 0.1% Triton X-100 in PBS for 1 h at room temperature. For nuclear staining, cells were permeabilized with 0.25% Triton X-100 in PBS for 10 min at room temperature and washed three times with PBS prior to blocking. For BrdU staining cells were incubated with 2M HCl, 0.1% Triton X-100 at 37°C for 10 min and then with 0.1M Sodium Borate (pH:8.5) for 30 min at room temperature followed by washing three times with PBS prior to blocking. Next, cells were incubated with primary antibodies, diluted in 1% NDS, 0.05% Triton X-100 in PBS overnight at 4°C. The next day, cells were washed three times with PBS and incubated with secondary antibodies diluted in 1% NDS, 0.05% Triton X-100 in PBS for 2h at room temperature. The nuclei of the cells were stained with ProLong Gold Antifade Reagent with DAPI (Cell Signaling). The following primary antibodies were used in this study: rabbit anti-GFAP (DAKO, 1:600), mouse anti-GFAP (Cell Signaling, 1:600), mouse anti-Tuj1 (Covance, 1:600), chicken anti-Tuj1 (Millipore, 1:1000), rabbit anti-Iba1 (WAKO, 1:400), rabbit anti-PDGFRa (Cell Signaling, 1:1000), mouse anti-NG2 (Millipore, 1:500), rat anti-BrdU (Oxford Biotech, 1:50), mouse anti-MAP2 (Millipore, 1:200), rabbit anti-Synapsin1 (Abcam, 1:200), rat anti-Mash1 (R&D Systems, 1:100), rabbit anti-Tbr2 (Abcam, 1:200), rabbit anti-Gsx2 (Millipore, 1:400), rabbit anti-Tox (Atlas antibodies, 1:200), mouse anti-vGlut1 (Millipore, 1:1000) and rabbit anti-GABA (Sigma, 1:10,000). The secondary antibodies used in this study were Alexa Fluor 546-, Alexa Fluor 488- and Alexa Fluor 647-conjugated secondary antibodies (Life Technologies). Images were acquired with a 20x or 40x objective (1024×1024 pixels, 1μm Z-step) using a Leica TCS SP8 confocal microscope (LEICA Microsystems). For each experiment measurements from 20-25 fields per coverslip were obtained for each condition.

### Electrophysiology

For whole-cell recordings iNs plated in PLO-laminin coated coverslips were used for electrophysiological experiments beginning at d15 up to d27 of the reprogramming protocol. The coverslips with the cells were placed onto a recording chamber and viewed using an Olympus CKX41 microscope with a 40x lens. The cells were bathed in a solution containing: 140mM NaCl, 2.8mMKCl, 2mM CaCl_2_, 4mM MgCl_2_, 10mM Glucose, and 20mM HEPES. For whole-cell recordings we used a capillary glass with Filament (Sutter instrument) to fabricate low resistance recording pipettes (∼5 MΩ) and filled with: 140mMKCl, 2mM CaCl_2_, 2mM MgCl_2_, 2mM Mg-ATP, 5mM EGTA and 10mMHEPES. Osmolarity and pH of all solutions were adjusted appropriately before experiments. Data were acquired at room temperature (22– 24°C) using an EPC9 HEKA amplifier and an ITC-16 acquisition system with a patch master software (HEKA). Data analysis was carried out using OriginPro8. **Voltage protocols:** The membrane of the cells was held at a holding potential of −70 mV and step depolarizing pulses were applied. Depolarization steps were applied for 50 msec in 10 mV increments from −80 mV to +50 mV with a sweep interval time of 1 sec and sweep duration of 500 ms.Each depolarizing pulse was proceeded by a hyperpolarizing step to −120 mV. **Current protocols:** Cells we held at their resting membrane potential (0pA) and depolarizing current steps from −20 pA to 200 pA from a holding current of 0pA were applied.

### RNA-Seq experiment and Bioinformatics analysis

For the RNA-Seq experiment, the following samples were prepared in 3 biological replicates: astrocytes (d1), sc-miRNA-transfected astrocytes (d7), miR-124-iNs (d7) and miR-124+ISX9-iNs (d7). Total RNA was extracted using the Nucleospin miRNA kit (Macherey-Nagel) according to manufacturer’s instructions. Libraries were prepared with TruSeq RNA Library Prep Kit v2 (Illumina) and 75c single-end sequencing in an Illumina NextSeq 550 sequencer. Raw libraries were quality checked and preprocessed using Fast QC (https://www.bioinformatics.babraham.ac.uk/projects/fastqc/). Mapping of reads against the Mouse Transcriptome (v.GRCm38.rel79) and transcript abundance estimation on Transcripts Per Million (TPM) values was performed using kallisto (Bray et al. 2016). Analysis of differential expression, interpretation and visualization was subsequently performed using kallisto-compatible Sleuth tool (Pimentel et al. 2017) and R-base functions. Gene ontology (GO) enrichment analysis was performed using the Gene Ontology Panther Classification System (http://pantherdb.org/).

### Analysis of AGO-CLIP-Seq data

AGO-HITS-CLIP datasets, performed in mouse brain cortex tissue (P13 neocortex, 5 replicates) and human brain tissues (motor cortex, cingulate gyrus) from 2 individuals, were retrieved from the publications (Chi et al. 2009) and (Boudreau et al. 2014) respectively. Raw libraries were quality checked using FastQC (www.bioinformatics.babraham.ac.uk/projects/fastqc/), while adapters/contaminants were detected utilizing an in-house developed pipeline and the Kraken suite (Davis et al. 2013). Pre-processing was performed with Trim Galore (Krueger 2015) and Cutadapt (Martin 2011). CLIP-Seq libraries were aligned against the reference genomes, i.e. GRCh38 and mm10 assemblies for human and mouse respectively, with GMAP/GSNAP (T. D. Wu and Nacu 2010) spliced aligner, allowing up to 2 mismatches. microCLIP CLIP-Seq-guided model (Paraskevopoulou et al. 2018) was utilized to identify binding events for the expressed miRNAs. In case of multiple replicates (i.e. mouse brain cortex) a miRNA binding event had to be present in at least two replicates to be considered as valid. Top expressed miRNAs were retrieved from the relevant publications. Human and mouse transcriptomes were compiled from ENSEMBL v96 (Cunningham et al. 2019) to annotate the retrieved miRNA binding events. Identified miRNA binding sites residing on 3’ UTR regions were retained and subsequently filtered to preserve only genes expressed in astrocytes. A list of ∼10,000 genes, expressed in astrocytes, with FPKM≥2, was retrieved from a reference publication and retained for analysis (Y. Zhang et al. 2014).

### 3’-UTR cloning and measurement of Luciferase activity

A cDNA fragment of 710 bp corresponding to the 3’-UTR of Zfp36l1, containing the 7-mer binding site of mmu-miR-124-3p.1 (5’-TACAGAAGCAACTTGA**<u>GTGCCTT</u>**-3’), was obtained from RNA that was isolated from astrocytes using oligo dT primer for cDNA synthesis and subsequently the specific primers **Zfp36l1FOR**: 5’-ATC*GAGCTC*CACATAAGGACAAGTCAATTT and **Zfp36l1REV**: 5’-TGC*TCTAGA*AGCTTTTCTCTCATTTGTTGTCA for PCR amplification. The PCR fragment was subcloned into pmiR-GLO reporter vector (Promega) at the *SacI* and *XbaI* restriction sites. In addition, a cDNA fragment of 860bp corresponding to the 3’UTR of Zfp36l2, containing the two adjacent putative 7-mer binding sites of mmu-miR-124-3p.1 (5’-GGGGAAATGGTCTCA**<u>GTGCCTT</u>**-3’ and 5’-CATAGGGCCCGAACT**<u>TGCCTTA</u>**-3’ respectively), was amplified as previously using the specific primers **Zfp36l2FOR**: 5’-ATC*GAGCTC*GTTCTTTTCACAGTAATATATGC-3’ and **Zfp36l2REV**: 5’-TG*CTCTAGA*CCAAAAAATTTTATTGGGGGAAAC-3’ respectively. All PCR products were subcloned into pmiR-GLO vector at the *SacI* and *XbaI* restriction sites and then were subjected to sequencing. Mutation of the verified 7-mer binding site of the 3’-UTR of Zfp36l1 was performed using the Q5-site directed mutagenesis kit (New England BioLabs) using the following primers: **Zfp36L1mutFOR**: 5’-TTTGTAATCTAACTTTGTCACTG-3’ and **Zfp36L1mutREV**: 5’-AAAATAAGTTGCTTCTGTAAACG-3’ and resulted in a mutated 7-mer binding site 5’-TACAGAAGCAACTTGA**TTTTTTT**-3’. All mutated clones were subjected to sequencing.

Luciferase assays were performed in HEK293T cells at 50-60% confluence. HEK293T cells were co-transfected with the 3’UTR-containing pmiR-GLO reporter constructs (1 μg) along with sc-miRNA or miR-124 mimics (80 nM, Thermo) using Lipofectamine 2000 (Thermo) and 48 h later luciferase activity was measured in cell lysates using the Firefly & Renilla Single Tube Luciferase Assay Kit (Biotium), according to manufacturer’s instructions. Firefly luciferase activity was normalized with Renilla luciferase activity measured in the same tube and the normalized Firefly luciferase activity for cells transfected with sc-miRNA was set to 1, thus estimating the fold change of normalized Firefly luciferase activity inflicted by miR-124 transfection for each construct.

### Target Site Blocker (TSB) experiment

For the functional validation of the miR-124/Zfp36l1 interaction a custom made miRCURY locked nucleic acid (LNA) miRNA Power Target Site Blocker (TSB) (Qiagen) was used with the following sequence: TTACAAGGCACTAAGTTGCTT. TSB was transfected in astrocytes along with sc-miRNA or miR-124-3p mimics using Lipofectamine 2000 (Invitrogen) in different molecular ratios: miR-124:TSB, 4:1 (80nM:20nM), 2:1 (80nM:40nM) and 1:1 (80nM:80nM).

### Western blot

Cells were washed once with ice-cold PBS and lysed for 15min in ice-cold lysis buffer (150mM NaCl, 50mM Tris (pH 7.5), 1%v/v Triton X-100, 1mM EDTA, 1mM EGTA, 0.1% SDS, 0.5% sodium deoxycholate) containing PhosSTOP phosphatase inhibitors and a complete protease inhibitor mixture (Roche Life Science), then centrifuged at 20,000 g for 15min, followed by collection of the supernatant and measurement of the protein concentration by Bradford assay (Applichem). Proteins were separated by SDS-polyacrylamide gel electrophoresis (PAGE) and transferred onto nitrocellulose membranes (Maine Manufacturing). Nonspecific binding sites were blocked in TBS, 0.1% Tween 20, 5% skimmed milk for 1 h at room temperature followed by overnight incubation with primary antibodies diluted in TBS, 0.1% Tween20, 5% BSA. Primary antibodies used were rabbit anti-Zfp36l1/Zfp36l2 (this antibody recognizes Zfp36l1 at 37kDa and Zfp36l2 at 51kDa) (Abcam, 1:500) and mouse anti-βactin (Millipore, 1:1000). Incubation with HRP-conjugated secondary antibodies, anti-mouse-HRP (Thermo, 1:10,000) and anti-rabbit-HRP (Thermo, 1:5,000) was performed for 2 h at room temperature and protein bands were visualized using the Clarity Western ECL Substrate (BIO-RAD).

### Lentiviral production

For lentiviral *in vivo* transduction, VSV-G (Vesicular Stomatitis Virus–Glycoprotein)– pseudotyped lentiviruses were used either for the over-expression of miR-124 along with GFP or as control expressing only GFP. More specifically, for lentiviral particles’ production, HEK 293T cells cultured in 10-cm Petri dishes at a 50-60% confluence were co-transfected with 10 μg lentiviral plasmid expressing miR-124-1 precursor under the CMV promoter and GFP under the EF1 promoter (SBI System Biosciences) or 10 μg lentiviral plasmid expressing GFP under the CMV promoter and the packaging plasmids pVSV-G (3.5 μg), MDL (6.5 μg), and RSV-REV (2.5 μg) (all kindly provided by Dr. Matsas’ lab) with calcium phosphate. The following day the culture medium was replaced with fresh one, the supernatant containing the lentiviral particles was collected 48 h and 72 h (second harvest) after transfection and concentrated by ultracentrifugation at 25,000 rpm (80,000 x g) for 2 h at 4°C using a sucrose gradient.

### *In vivo* reprogramming protocols

This study was carried out in strict compliance with the European Directive 2010/63/EU and the Greek National Law 161/91 for Use of Laboratory Animals, according to FELASA recommendations for euthanasia and the National Institutes of Health Guide for Care and Use of Laboratory Animals. All protocols were approved by the Animal Care and Use Committee of the Hellenic Pasteur Institute (Animal House Establishment Code: EL 25 BIO 013). License No 2585/31-5-18 and 490073/15-6-21 for the experiments was issued by the Greek authorities of the Veterinary Department of the Athens Prefecture. The manuscript was prepared in compliance with the ARRIVE guidelines for reporting animal research.

### Cortical Trauma model

Adult male and female FVB or Rosa26-EYFP reporter mice (8-16 weeks old) were deeply anaesthetized using inhalable isoflurane, and positioned in a stereotaxic apparatus. The dorsal surface of the skull was exposed through a midline incision and a burr hole was drilled at the following coordinates: antero-posterior (AP) −1.0 mm, caudal to Bregma; lateral (L) 1.0 mm to the midline. A 26-gauge needle was inserted into the brain parenchyma in a depth of 0.9 mm from the surface of the brain to create a trauma in the cortex, avoiding the corpus callosum and hippocampus. The inserted needle was moved along the anterior-posterior axis between positions (AP) −1.1 and −0.9 to widen the trauma. The skin was sutured, a local analgesic cream containing 2.5% lidocain and 2.5% prilocain was applied and the animals were kept warm until they were fully awake. Viral injection took place 4 days after the cortical trauma.

### Lentiviral injection in the cortex of FVB mice

A 10μl Hamilton syringe (Hamilton) with a 26-gauge needle was slowly inserted into the brain tissue at coordinates (AP) −1.1 mm, (L) 1.0 mm, and (V): 1.0 mm, from the same burr hole on the skull and 2μl of lentiviral concentrate was injected at a rate of 0.5 μl/min. The needle was left in position for 5 min after each injection and then withdrawn gently. A second viral injection was repeated at coordinates (AP) −0.9 mm, (L) 1.0 mm, and (V): 1.0 mm with similar procedures, and surgery was completed as described above. A group of animals was injected with the lentivirus LV-miR-124-GFP and another one with the control lentivirus LV-GFP. A subgroup of the LV-124 group received intraperitoneally 20 mg/kg of ISX9 (Tocris) diluted in (2-Hydroxypropyl)-β-cyclodextrin (Sigma) (ISX9 concentration: 2 mg/ml in 30% (2-Hydroxypropyl)-β-cyclodextrin (Sigma) diluted in sterile ddH_2_O) (Petrik et al. 2012; Kutsche et al. 2018) once a day, for 5 consecutive days, beginning 48 h p.i. Animals were sacrificed 7 days or 3 weeks after viral injection.

### AAV injection in the cortex of R26R-EYFP mice

1,5μl of a viral mix (1:1) of AAV9:CMV-miR-124-mCherry (titer: ≥5×10^12^ GC/ml, GeneCopoeia) and AAV5:GFAP-Cre (titer:≥7×10¹² GC/ml, Addgene) viruses were injected with a 10μl Neuros syringe (Hamilton) with a 33-gauge blunt needle, at a rate of 0.25 μl/min, as described above. One subgroup in each time point (d7, 3w and 8w) received i.p. 20 mg/kg of ISX9 as described above. BrdU (Sigma) (50mg/kg) was administered twice a day, beginning at d2 after trauma for 4 consecutive days.

### Tissue Preparation, Histology, and Immunohistochemistry

For histology, mice were deeply anaesthetized by inhaling isoflurane, and perfused with 4% paraformaldehyde (PFA) via left cardiac ventricle. The brains were removed, post-fixed in 4% PFA overnight and then cryo-protected in 20% sucrose overnight. Tissues were then frozen in −20°C isopentane and cut into 20 μm-thick coronal sections on a cryostat (Leica CM1900), collected on silane-coated slides and stored at −20°C. For detection of specific antigens with immunofluorescence, sections were left for 15 min in room temperature, washed in PBS, and blocked with 5% normal goat or donkey serum (Merck-Millipore) in PBS-T (0.1% Triton X-100 in PBS) for 1 h. Incubation with primary antibodies took place overnight at 4°C. Primary antibodies used were: chicken polyclonal anti-GFP (Abcam, 1:1000), mouse monoclonal anti-neuronal nuclei (NeuN) (Merck-Millipore, 1:300), rabbit polyclonal anti-glial fibrillary acidic protein (GFAP) (Dako,1:600), mouse monoclonal anti-GFAP (Cell Signaling, 1:600); rabbit polyclonal anti-oligodendrocyte transcription factor 2 (Olig2) (Merck-Millipore, 1:200), rabbit polyclonal anti-ionized calcium-binding adapter 1 (Iba-1) (Wako, 1:600) rabbit polyclonal anti-Tbr1 (Abcam,1:250), rat monoclonal anti-BrdU (Oxford Biotech, 1:100), rat monoclonal anti-red fluorescent protein (RFP) (Chromotek, 1:1000) and goat anti-mCherry (1:1000). Following incubation with primary antibodies, sections were washed with PBS and incubated for 2 h with the appropriate secondary antibodies conjugated with Alexa Fluor 488 (green), 546 (red), or 647 (blue). For nuclei staining sections were incubated with Hoechst (Molecular Probes) and coverslipped with Mowiol (Calbiochem) or with DAPI using the Everbrite mounting medium with DAPI (Biotium). Images were acquired with a 40x objective using Leica TCS SP8 and Leica TCS-SP5II confocal microscopes (LEICA Microsystems).

### Image analysis

Images were analyzed using Fiji/ImageJ software (National Institutes of Health) and Imaris Software (Bitplane).

**In vitro analysis:** mean fluorescence intensity of Tbr2, Tox and Mash1 staining inside the cell nuclei was quantified using a custom-written macro implemented in Fiji. Initially automatic detection of nuclei was performed using the Otsu method and Tuj1+cells were selected based on their mean intensity value above a user-defined threshold of 40, followed by a manual validation according to cell morphology (cells with an astrocyte-like morphology with a big, rectangular soma and none or few processes were excluded). Mean fluorescence intensity of Tbr2, Tox and Mash1 inside the cell nuclei ROIs was measured both forTuj1- and Tuj1+ cells. Quantification was performed in maximum intensity projections. For each experiment measurements from at least 200-300 cells were obtained for each condition.

Morphological characterization of Tuj1+ cells in the TSB blocker experiments was also conducted using Fiji. More specifically, morphological characterization included quantification of the cell body area and the number of processes extending from the soma of Tuj1+ cells. Tuj1+ cells were selected based on the mean intensity value of their soma above a user-defined threshold and were sorted in 3 groups: cells with a multipolar morphology bearing 3 or more processes extending from the soma, cells with 1-2 processes and cells exhibiting an astrocyte-like morphology with a rectangular soma possessing none or 1-2 processes. Quantification was performed in maximum intensity projections. For each experiment measurements from at least 100-150 cells were obtained for each condition.

**In vivo analysis:** for each animal and each immunofluorescence staining, cell counting was performed on brain coronal sections collected at 240 mm intervals across the whole antero-posterior extent of the hippocampus (Bregma −0.5mm up to −2.5mm) in a total number of 3-4 mice for each experimental condition. For estimation of the percentage of LV or AAV transduced cells that have a specific phenotype, images of GFP+ or mCherry+ cells respectively found in each set of sections were acquired and double- or triple-positivity with cell type-specific markers was evaluated. All GFP+ or mCherry+ cells found in the cortex within these sections were imaged and analyzed. For the analysis of AAV9-transduced cells mCherry+ mature neurons were excluded from the analysis. To isolate and evaluate morphological features of astrocytes, resident neurons and iNs we used Imaris Contour Surface and Surface Segmentation (Imaris v.9.3.1, Bitplane) for each channel represented in **Suppl Fig. 9**. Same threshold values were used for all groups during the analysis.

### Statistical analysis

All *in vitro* quantified data are presented as average ± SD, unless otherwise indicated. Two-tailed Student t-test was used to calculate statistical significance with p values for all the data obtained from the experiments with the TSB blocker, while for the rest of the data a one-tailed Student t-test was used. p values less than 0.05 (p<0.05) were considered indicative of significance. *In vivo* data were assessed using a one-way analysis of variance (ANOVA). When interactions were detected, group comparisons were performed using a two-sample assuming unequal variances test.

## Acknowledgments

This work was financially supported by: ‘BIOIMAGING-GR: A Greek Research Infrastructure for Visualizing and Monitoring Fundamental Biological Processes (MIS 5002755)’, funded by the Operational Program “Competitiveness, Entrepreneurship and Innovation” (NSRF 2014-2020), co-financed by Greece and the European Union (European Regional Development Fund); ARISTEIA-II *‘*Astro-Rep’ 3713 Excellence Grant of the Greek Ministry of Education and Fondation Santé Grant 2017-2018, awarded to DT. We also acknowledge funding from the Stavros Niarhos Foundation (SFN) Grant to the Hellenic Pasteur Institute, as part of the Foundation’s initiative to support the Greek Research Center Ecosystem; Greek General Secretariat of Research and Technology ‘Action for the Study of Neurodegenerative Diseases on the Basis of Precision Medicine’ and ‘KRIPIS-II’ Action (MIS 5002486) under the Operational Strategic Reference Framework 2014–2020. We would like to thank Dr. Era Taoufik for critical comments on the manuscript.

## Author Contributions

EP and DT conceived whole project, designed experiments and analyzed the data; EP, CG, MG and MM conducted in vitro experiments; DCT and SJT designed and performed electrophysiology experiments; TK, DK and AGH designed and performed bioinformatics analysis and EP contributed in analyzing the data; PNK and IT designed and performed *in vivo* experiments, PKN and EP analyzed relevant data and PKN contributed in paper writing; EX developed the Fiji macro and helped with image analysis workflow; EP and DT wrote the manuscript; DT supervised the project and acquired funding.

## Data Availability

High throughput RNA-Sequencing data are deposited in the European Nucleotide Archive under Study accession PRJEB38603.

