## Supplemental information for "A miR-124-mediated post-transcriptional mechanism controlling the cell fate switch of astrocytes to induced-neurons"

### Suppl. Figure 1

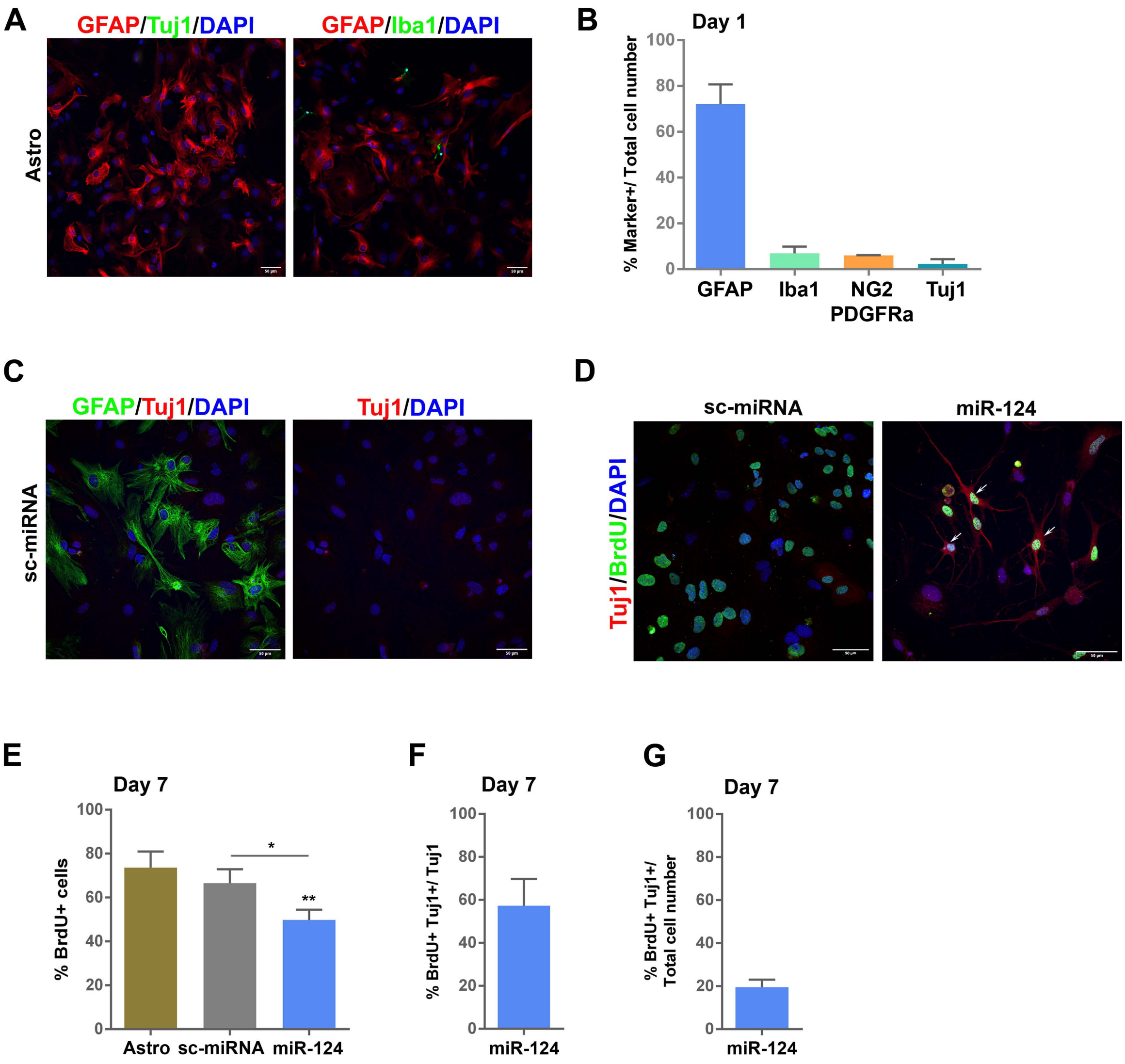

**Supplementary Figure 1**

**(A)** Co-immunostaining of the primary cortical astrocytes' culture with anti-GFAP/Tuj1 and GFAP/Iba1. **(B)** Quantification of the percentage of GFAP+ astrocytes, Iba1+ microglia, NG2+/PDGFRa+ OPCs and Tuj1+ neurons in the primary astrocytic culture, (average ± SD, n=3 independent experiments). **(C)** Co-immunostaining of astrocytes transfected with sc-miRNA at d7 of the reprogramming protocol with anti-GFAP/Tuj1 antibodies. **(D)** Co-immunostaining of astrocytes transfected with sc-miRNA or miR-124 and supplemented with 10μM BrdU for 4 consecutive days at d7 of the reprogramming protocol with anti-BrdU/Tuj1 antibodies. **(E)** Quantification of the percentage of BrdU+ cells in the control astrocytic culture or in astrocytes transfected with sc-miRNA or miR-124 at d7 of reprogramming (average ± SD, n=3 independent experiments, **p<0.01 *vs* astrocytes and *p<0.05 for miR-124 *vs* sc-miRNA). Quantification of the percentage of Tuj1+ reprogrammed iNs at d7 that originated from a proliferating state (BrdU+/Tuj1+ double positive) *vs* the total Tuj1+ reprogrammed iNs **(F),** or *vs* the total cells in culture **(G)**, (average ± SD, n=3 independent experiments).

### Suppl. Figure 2

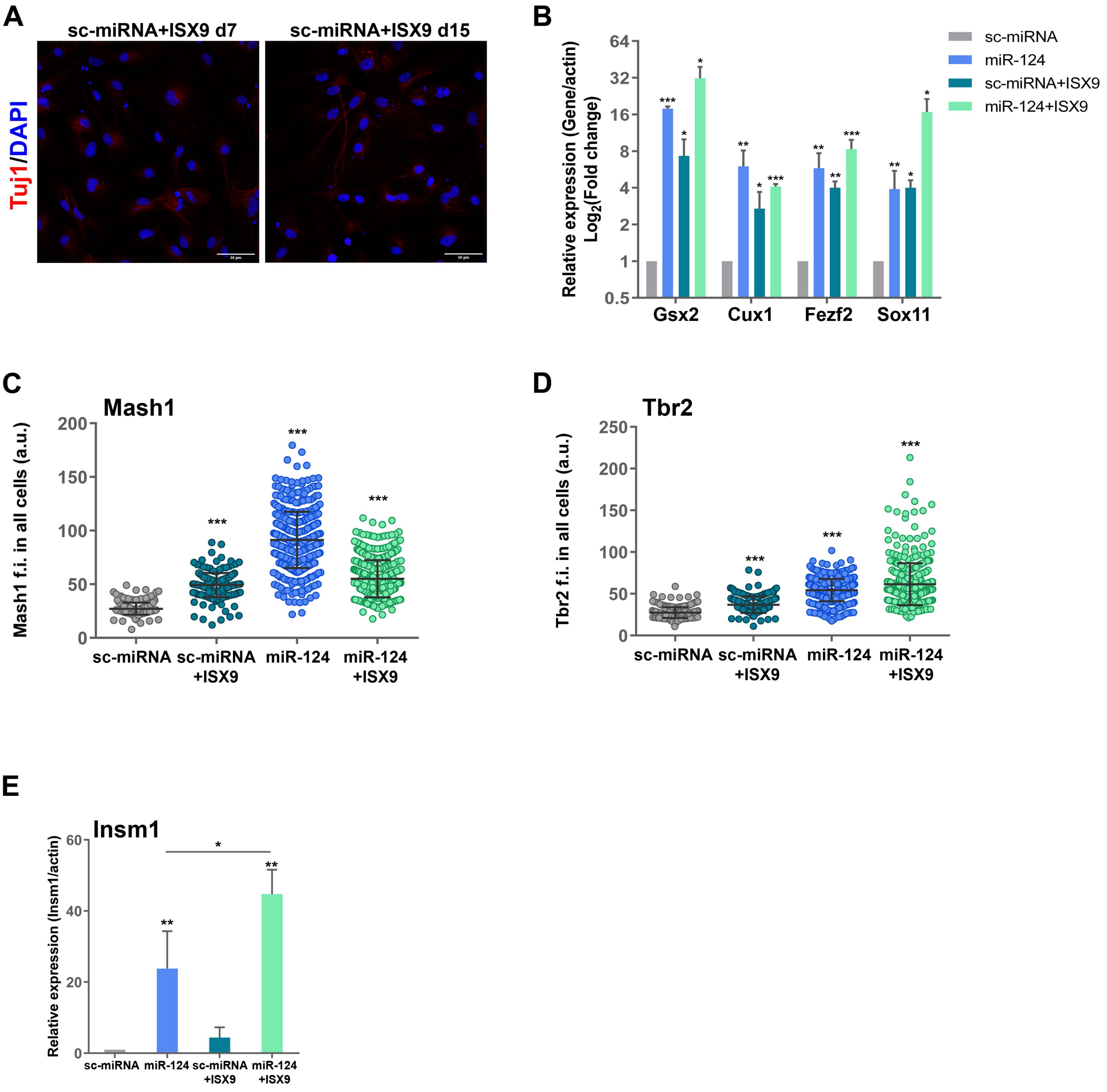

**Supplementary Figure 2**

**(A)** Immunostaining of astrocytes transfected with sc-miRNA supplemented with ISX9 at d7 and d14 of the reprogramming protocol with anti-Tuj1 antibody. **(B)** RT-qPCR analysis of the mRNA levels of *Gsx2, Cux2, Fezf2* and *Sox11* at d7 of the reprogramming protocol. Data are presented as log$_2$(fold change) *vs* sc-miRNA (average ± SD, n=3 independent experiments, *p<0.05, **p<0.01 and ***p<0.001 *vs* sc-miRNA). Measurement of the mean nuclear fluorescence intensity of Tbr2 **(C)** and Mash1 **(D)** in astrocytes transfected either with sc-miRNA or miR-124 in the presence or absence of ISX9 at d7 of the reprogramming protocol (co-immunostaining with anti-Tbr2/Mash1 antibodies). A representative experiment is shown of n=3 independent experiments (mean ± SD, n=250 cells for sc-miRNA, n=220 cells for sc-miRNA+ISX9, n=490 for miR-124 and n=450 cells for miR-124+ISX9, ***p<0.001 *vs* sc-miRNA). (**E**) RT-qPCR analysis of the mRNA levels of the TF and intermediate progenitors' (IP) marker *Insm1* at d7 (average ± SD, n=3 independent experiments, **p<0.01 vs sc-miRNA, *p<0.05 *vs* miR-124).

### Suppl. Figure 3

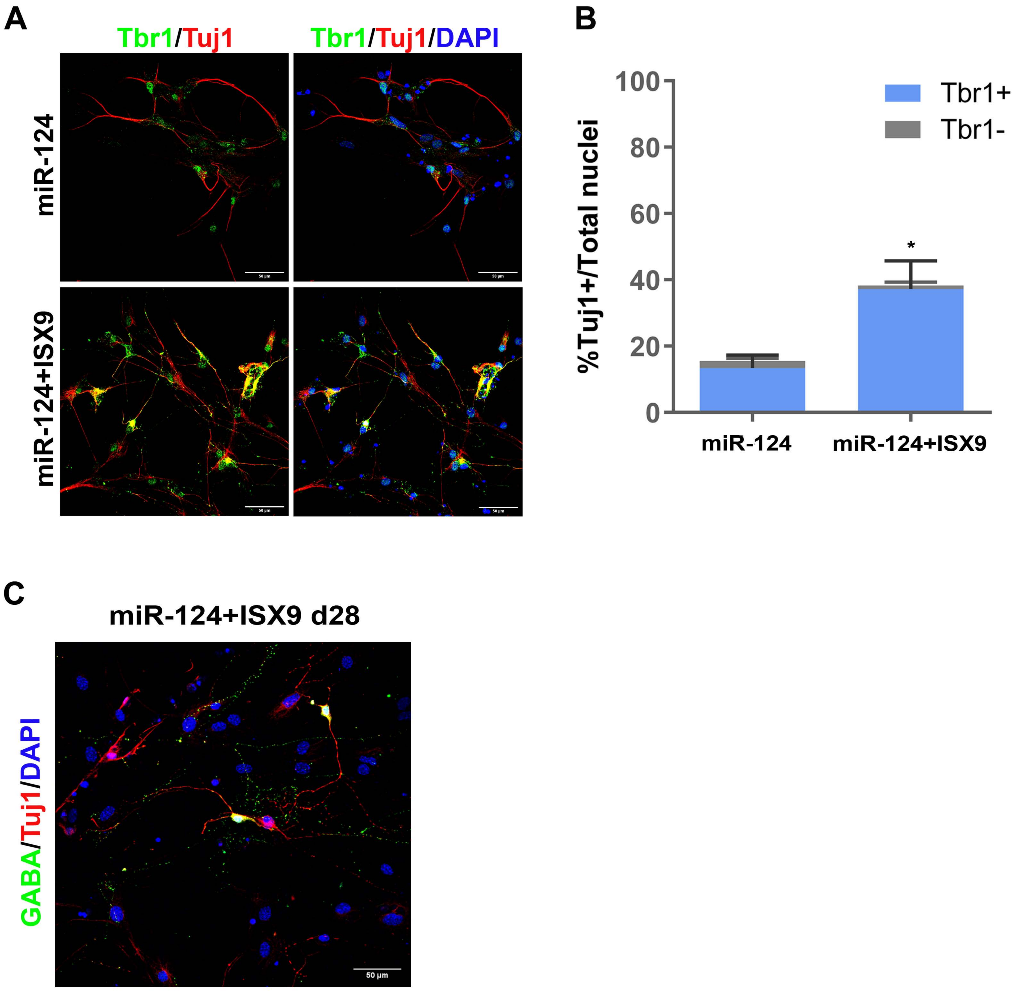

**Supplementary Figure 3**

**(A)** Co-immunostaining of miR124-iNs and miR-124+ISX9-iNs at d14 of the reprogramming protocol with anti-Tbr1/Tuj1 antibodies. **(B)** Quantification of the percentage of Tuj1+ iNs at d14 of reprogramming. The percentage of Tbr1+/Tuj1+ double positive iNs is shown in blue (average ± SD, n=3 independent experiments, *p<0.05 refers to %Tuj1+ miR-124+ISX9-iNs *vs* miR-124-iNs, no statistical significant difference was found for %Tbr1+/Tuj1+ double positive miR-124+ISX9-iNs *vs* miR-124-iNs). **(C)** Co-immunostaining of miR-124+ISX9-iNs at d28 of the reprogramming protocol with anti-GABA/Tuj1 antibodies.

### Suppl. Figure 4

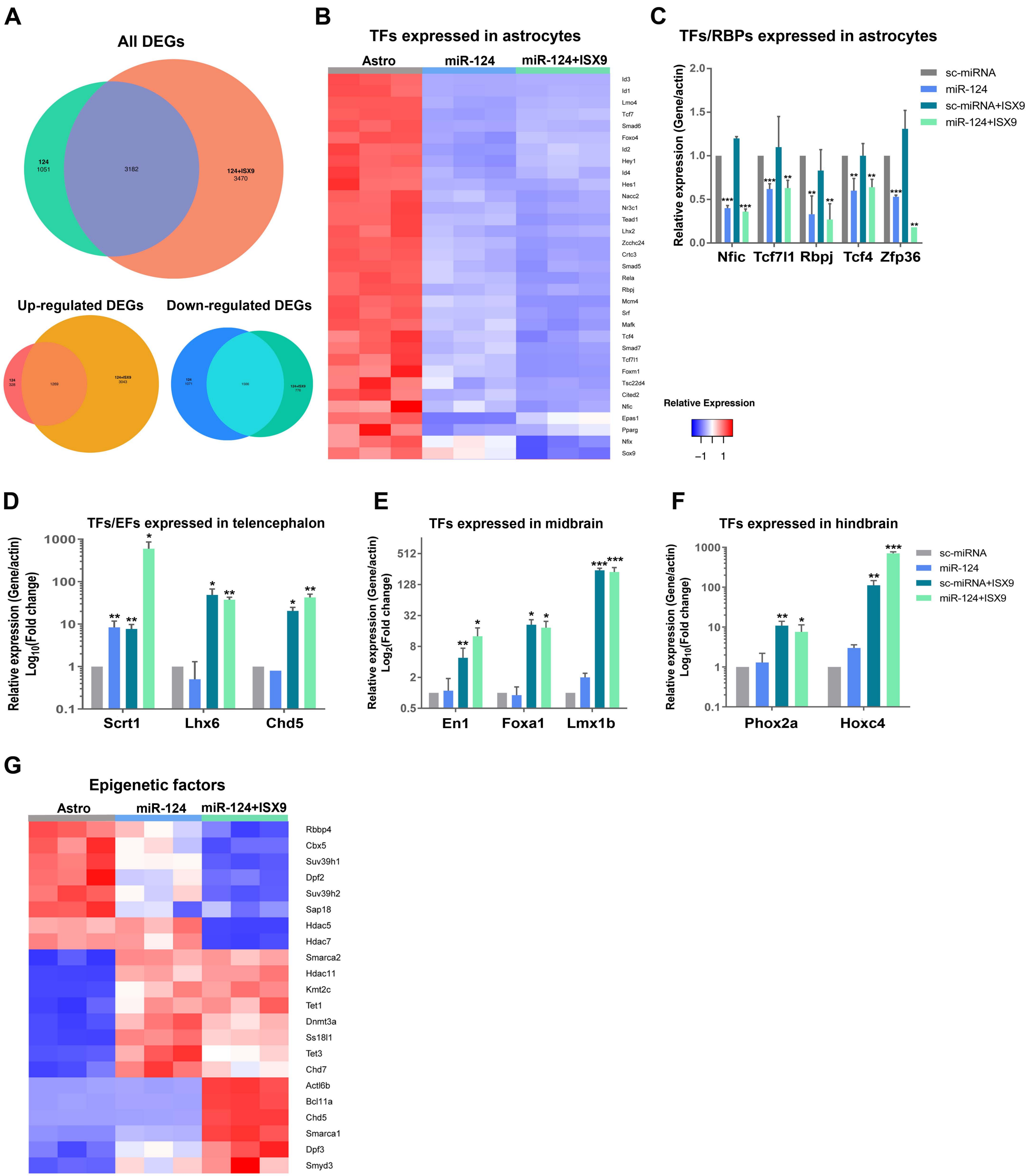

**Supplementary Figure 4**

**(A)** Venn diagrams presenting the common total (top), up-regulated (bottom left) and down-regulated (bottom right) DEGs between miR-124 *vs* astro and miR-124+ISX9 *vs* astro datasets ($1 \leq \log_2$(fold change)$\leq -1$, FDR<0.05). **(B)** Heat map analysis of 35 down-regulated astrocytic TFs. **(C)** RT-qPCR validation of the mRNA levels of the TFs *Nfic, Tcf7l1, Rbpj, Tcf4* and the RBP *Zfp36* expressed in astrocytes. Data are presented as fold change *vs* sc-miRNA (average ± SD, n=3 independent experiments, **p<0.01 and ***p<0.001 *vs* sc-miRNA). RT-qPCR validation of the mRNA levels of the TFs *Scrt1* and *Lhx6* and the epigenetic factor (EF) *Chd5* expressed in telencephalon **(D)**, the TFs *En1, Foxa1* and *Lmx1b* expressed in midbrain **(E)** and the TFs *Phox2a, Hoxc4* expressed in hindbrain **(F)** at d7 of the reprogramming protocol, (average ± SD, n=3 independent experiments, *p<0.05, **p<0.01, ***p<0.001 *vs* sc-miRNA). Data are presented as $\log_{10}$(fold change) **(D, F)** and $\log_2$(fold change) **(E)** *vs* sc-miRNA. **(G)** Heat map analysis of 22 up- and down-regulated epigenetic factors (EFs).

### Suppl. Figure 5

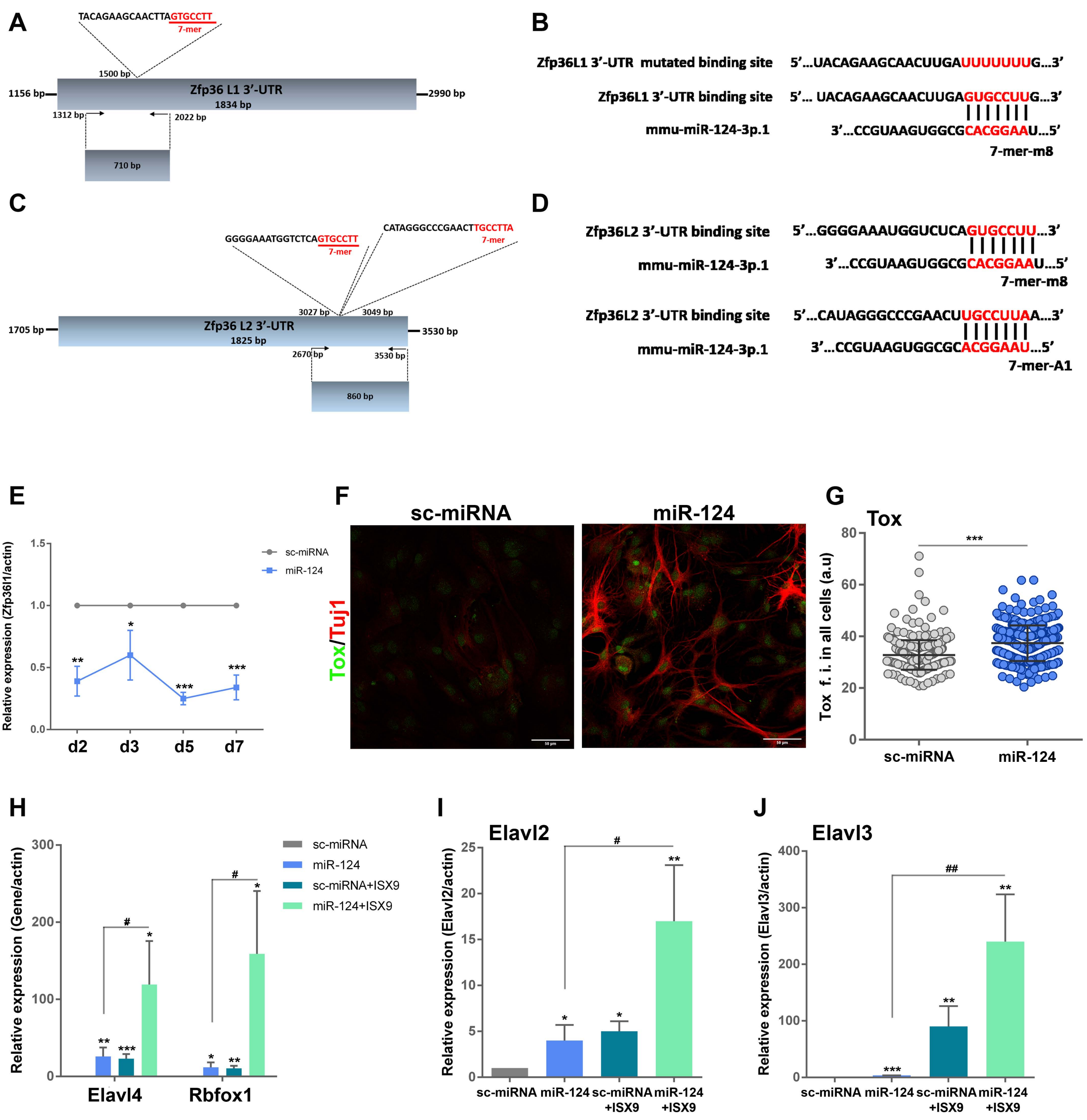

**Supplementary Figure 5**

**(A)** RT-qPCR analysis of the mRNA levels of *Zfp36l1* at the early time points d2, d3, d5 and d7 of the reprogramming protocol (average ± SD, n=3 independent experiments, *p<0.05, **p<0.01, ***p<0.001 *vs* sc-miRNA). **(B)** Co-immunostaining of astrocytes transfected with sc-miRNA or miR-124 at d5 with anti-Tox/Tuj1 antibodies. **(C)** Measurement of the mean nuclear fluorescence intensity of Tox in astrocytes transfected either with sc-miRNA or miR-124 at d5. A representative experiment is shown of n=2 independent experiments (mean ± SD, n=323 cells for sc-miRNA and n=357 cells for miR-124, ***p<0.001 *vs* sc-miRNA). RT-qPCR analysis of the mRNA levels of the Zfp36l1 targets *Elavl4* and *Rbfox1* **(D)** and the other members of the nElavl family *Elavl2* **(E)** and *Elavl3* **(F)** at d7 (average ± SD, n=3 independent experiments, *p<0.05, **p<0.01, ***p<0.001 *vs* sc-miRNA and $^{\#}$p<0.05, $^{\#\#}$p<0.01 *vs* miR-124).

### Suppl. Figure 6

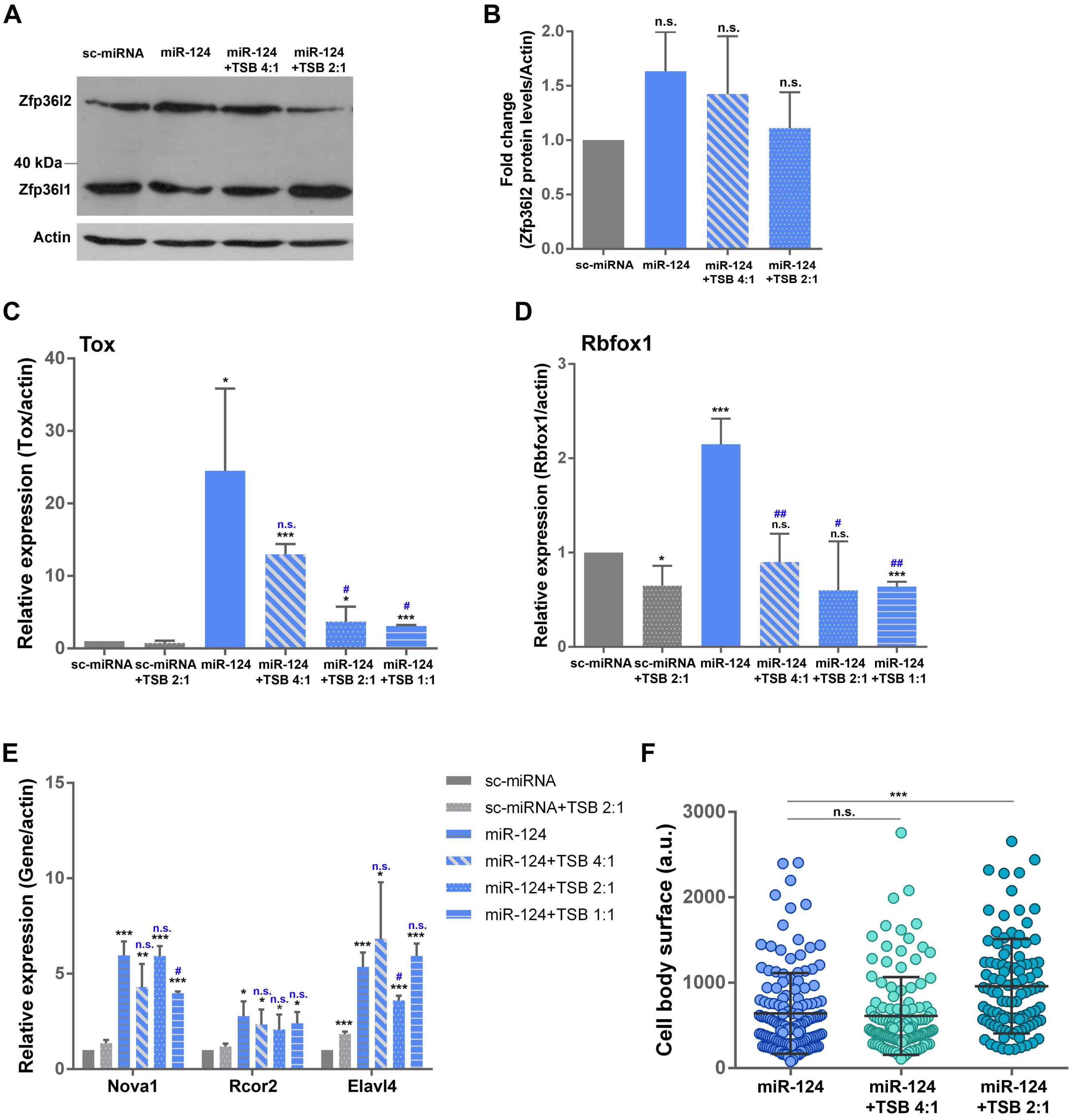

**Supplementary Figure 6**

**(A)** Western blot analysis of Zfp36l2 (51 kDa) and Zfp36l1 (37 kDa) protein levels in astrocytes transfected with sc-miRNA or miR-124 in the presence or absence of TSB (miR-124:TSB molecular ratio 4:1 and 2:1) at d5 of the reprogramming protocol. Actin has been used as loading control. **(B)** Quantification of Zfp36l2 protein levels and normalization with β-actin. Normalized Zfp36l2 levels for astrocytes transfected with sc-miRNA have been set to 1, (average ± SD, n=3 independent experiments). RT-qPCR analysis of the mRNA levels of the Zfp36l1 targets *Tox* **(C)**, *Rbfox1* **(D)** and *Nova*1, *Rcor2* and *Elavl4* **(E)** in the presence or absence of increasing concentrations of TSB (miR-124:TSB molecular ratio 4:1, 2:1 and 1:1) at d5 of the reprogramming process (average ± SD, n=3 independent experiments, *p<0.05, ***p<0.001 *vs* sc-miRNA and $^{\#}$p<0.05, $^{\#\#}$p<0.01 *vs* miR-124). **(F)** Measurement of the cell body surface of Tuj1+ cells. A representative experiment is shown of n=3 independent experiments (mean ± SD, n=141 for miR-124, n=121 for miR-124+TSB 4:1 and n=101 for miR-124+TSB 2:1, ***p<0.001 *vs* miR-124).

#### Suppl. Figure 7

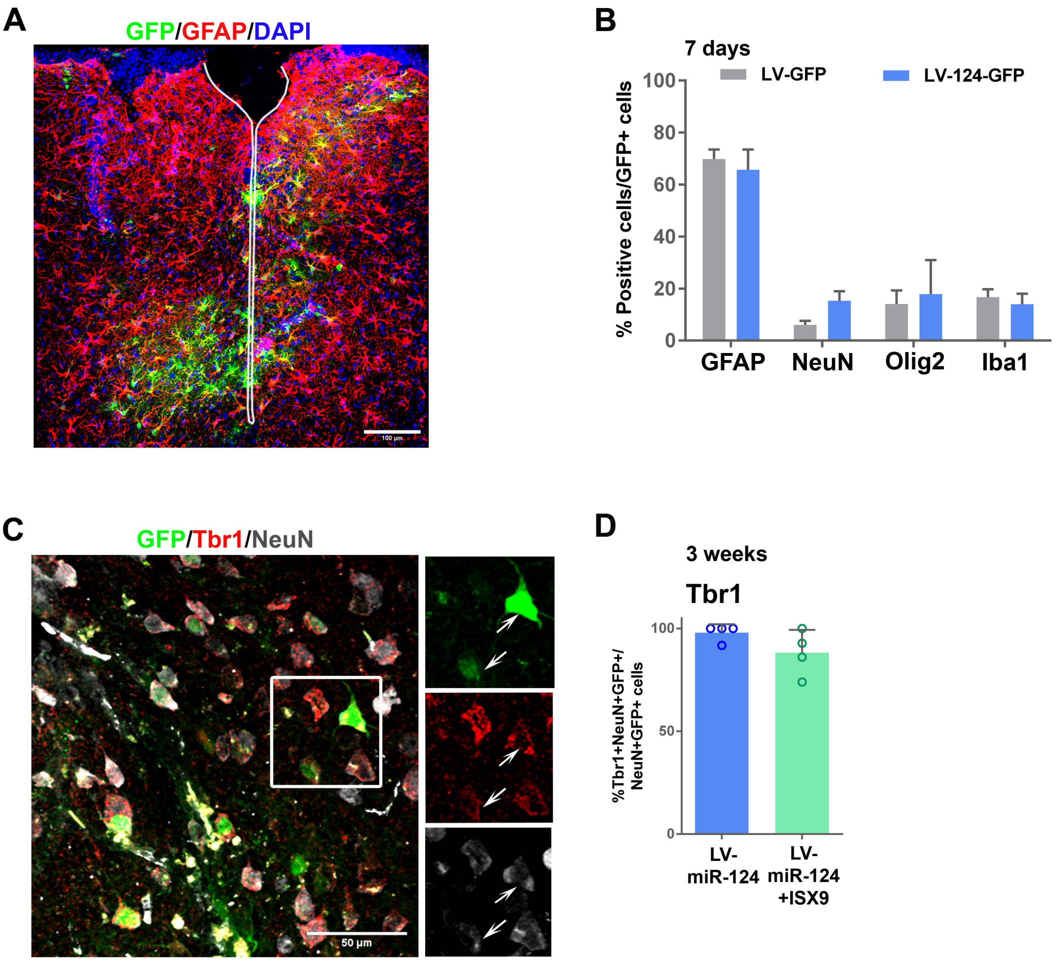

**Supplementary Figure 7**

**(A)** Image of astrogliosis (red, GFAP staining) surrounding cortical trauma (white continuous line), 7d p.i. of LV-124-GFP (transduced cells in green). **(B)** Brain cell types transduced by LV-GFP and LV-124-GFP, 7d p.i. Both lentiviruses preferably transduce GFAP+ astrocytes, and to a lesser extent NeuN+ neurons, Olig2+ oligodendrocytes, and Iba-1+ microglia, (average ± SD, n=4 animals per group per staining). **(C)** LV-miR-124-transduced cells in the peritraumatic cortical parenchyma expressing the cortical neuronal marker Tbr1, 3w p.i. Image from an animal transduced with LV-miR-124 and co-treated with ISX9. Inset area indicated in white frame, white arrows indicate NeuN+/Tbr1+ transduced cells. **(D)** Percentage of NeuN+ LV-124-transduced iNs also expressing Tbr1, 3w p.i. with or without treatment with ISX9, (average ± SD, n=4 animals for both groups).

#### Suppl. Figure 8

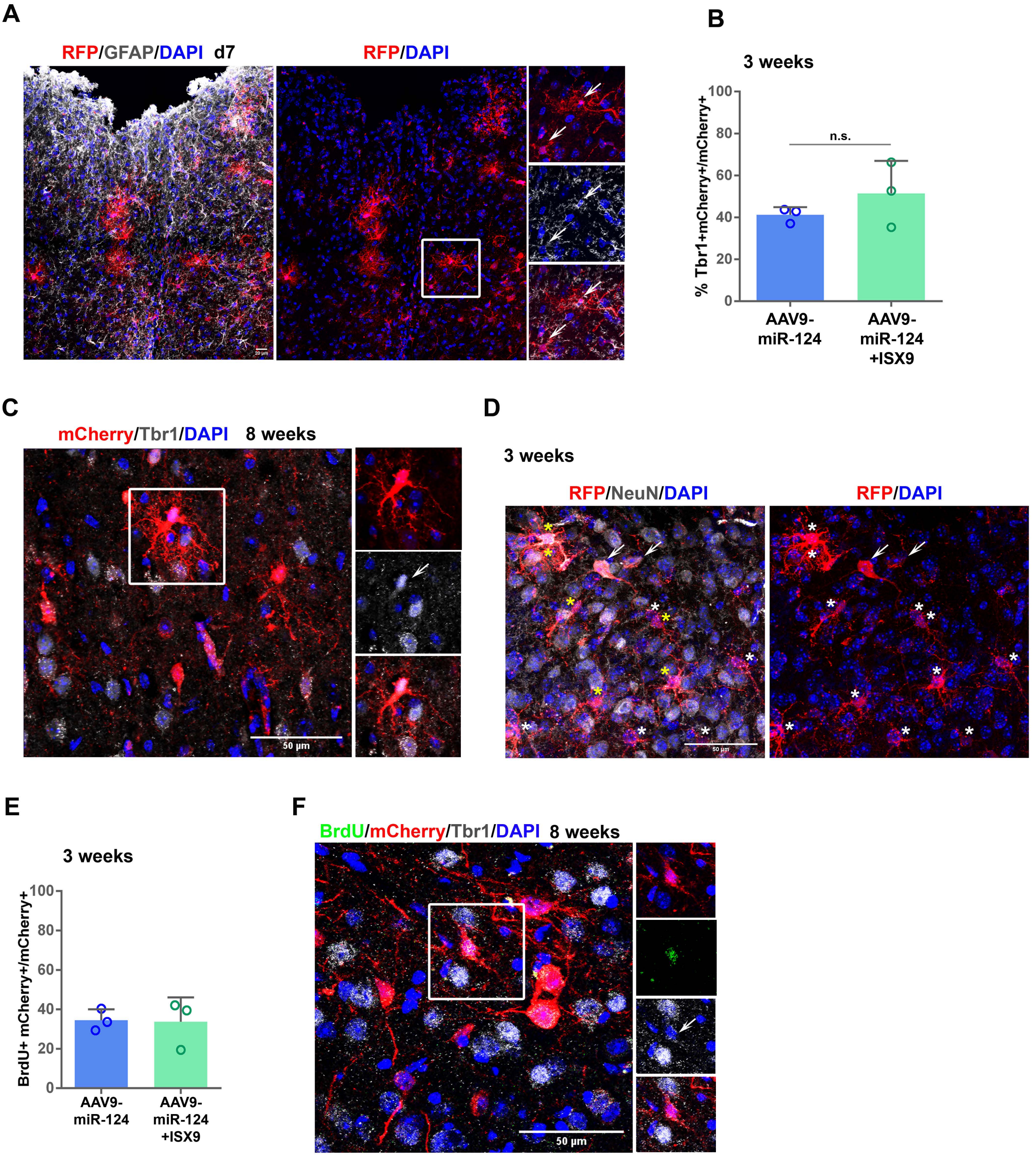

**Supplementary Figure 8**

**(A)** Image of astrogliosis (grey, GFAP staining) of the peritraumatic area, 7d p.i. of AAV9-124-mCherry (transduced cells in red detected with anti-RFP antibody). Inset area indicated in white frame, white arrows indicate GFAP+/RFP+ astrocytes. **(B)** Percentage of AAV9-124-transduced cells expressing the cortical marker Tbr1, 3w p.i. with or without treatment with ISX9, (average ± SD, n=3 animals for both groups). **(C)** AAV9-miR-124-transduced iNs in the peritraumatic cortical parenchyma 8w p.i. co-stained with anti-mCherry and anti-Tbr1 antibodies. Inset area indicated in white frame shows a representative mCherry+ transduced iN positive for the cortical neuronal marker Tbr1 (white arrow), still exhibiting a branched immature neuronal morphology. **(D)** AAV9-miR-124-transduced cells in the peritraumatic cortical parenchyma 3w p.i. co-stained with anti-mCherry and anti-NeuN antibodies, exhibiting an immature iN phenotype with a branched morphology (indicated in asterisks), some of them being mCherry+/NeuN+ (orange asterisks). Endogenous mCherry+/NeuN+ neurons distinguished by their bigger conical soma and bipolar morphology (arrows) were excluded from the analysis. **(E)** Percentage of AAV9-miR-124-transduced cells that have incorporated BrdU, 3w p.i. in the presence or absence of ISX9, (average ± SD, n=3 animals for both groups). **(F)** AAV9-miR-124-transduced cells in the peritraumatic cortical parenchyma 8w p.i. co-stained with anti-mCherry, anti-Tbr1 and anti-BrdU antibodies. Inset area indicated in white frame shows a representative Tbr1+/mCherry+ iN that had incorporated BrdU still viable after 8w p.i.

#### Suppl. Figure 9

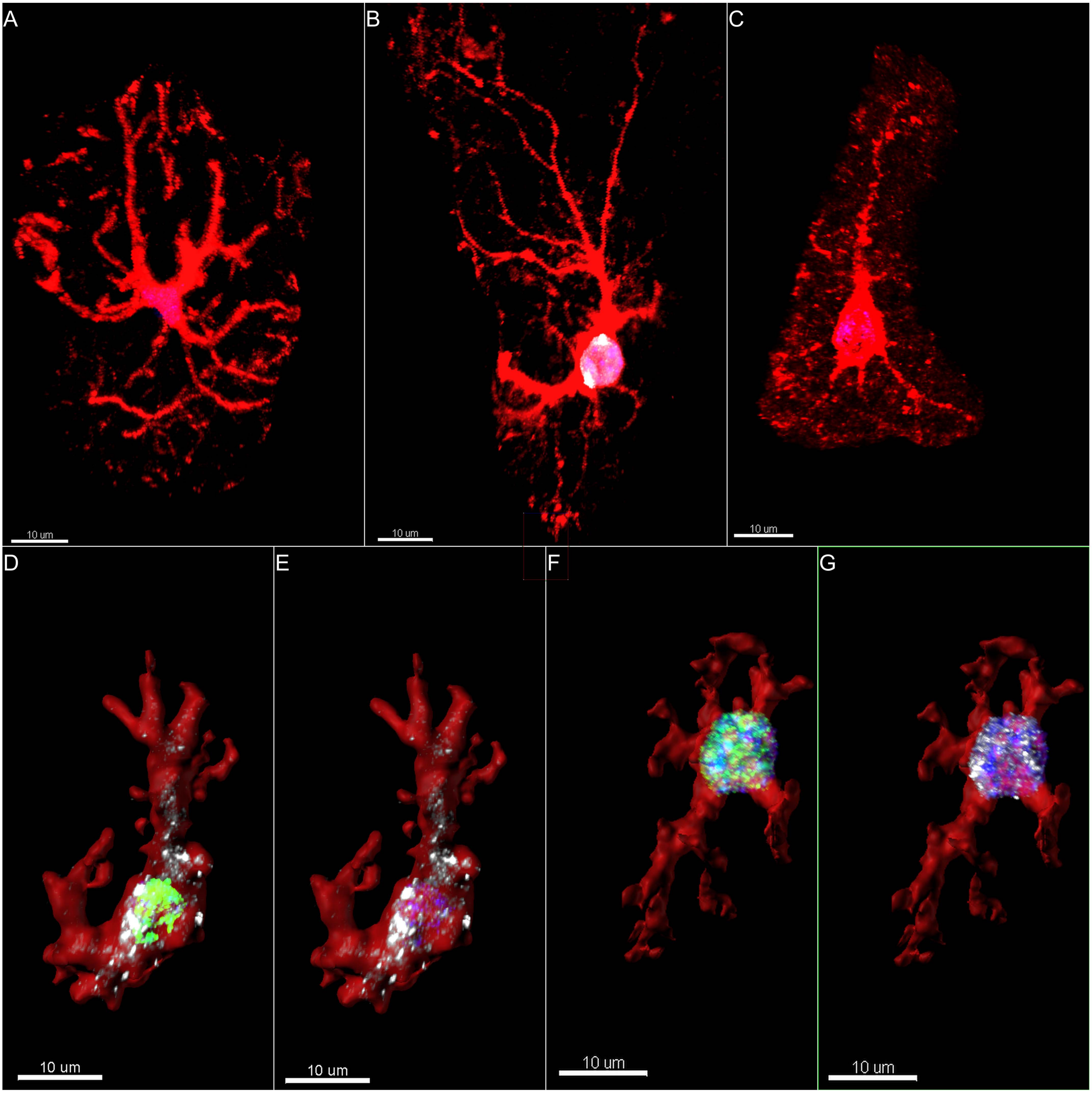

**Supplementary Figure 9**

Typical morphological features of transduced (mCherry+ in red) resident astrocytes **(A)**, NeuN+ iNs (white) exhibiting a transitional morphology 3w p.i. **(B)** and resident neurons **(C)**. Star-like astrocytic morphology is being replaced by an evident polarization in the soma and the processes of iNs, which can be still distinguished from mature resident neurons exhibiting defined localization of dendrites and the axon. Similar characteristic immature neuronal morphology is observed in iNs that have originated from proliferating cells 3w **(D-E)** and 8w p.i **(F-G)**. 3D representation of mCherry (red), BrdU (green) Tbr1 (white), DAPI (blue) labeling using Imaris Software.

**Supplementary Data 1 (Excel file):** Direct AGO-HITS-CLIP-derived miR-124-3p binding sites in 3' UTR regions in mouse brain cortex, characterized by microCLIP analysis and defined significantly down-regulated in miR-124 *vs* sc-miRNA RNA-Seq dataset (log2(fold change)≤-1, FDR<0.01).

**Supplementary Data 2 (Excel file):** Direct Zfp36l1-iCLIP-derived targets in T and B cells, defined significantly up-regulated in miR-124 *vs* sc-miRNA RNA-Seq dataset ($log_2$(fold change)≥1, FDR<0.05).

**Supplementary Data 3 (Excel file):** Differentially expressed astrocytic and neuronal genes presented in the heat map of **Fig 4A.**
